# Reduced oriens-lacunosum/moleculare (OLM) cell model identifies biophysical current balances for *in vivo* theta frequency spiking resonance

**DOI:** 10.1101/2022.10.20.513073

**Authors:** Zhenyang Sun, David Crompton, Milad Lankarany, Frances K Skinner

## Abstract

Conductance-based models have played an important role in the development of modern neuroscience. These mathematical models are powerful “tools” that enable theoretical explorations in experimentally untenable situations, and can lead to the development of novel hypotheses and predictions. With advances in cell imaging and computational power, multi-compartment models with morphological accuracy are becoming common practice. However, as more biological details are added, they make extensive explorations and analyses more challenging largely due to their huge computational expense. Here, we focus on oriens-lacunosum/moleculare (OLM) cell models. OLM cells can contribute to functionally relevant theta rhythms in the hippocampus by virtue of their ability to express spiking resonance at theta frequencies, but what characteristics underlie this is far from clear. We converted a previously developed detailed multi-compartment OLM cell model into a reduced single compartment model that retained biophysical fidelity with its underlying ion currents. We showed that the reduced OLM cell model can capture complex output that includes spiking resonance in *in vivo*-like scenarios as previously obtained with the multi-compartment model. Using the reduced model, we were able to greatly expand our *in vivo*-like scenarios. Applying spike-triggered average analyses, we were thus able to to determine that it is a combination of hyperpolarization-activated cation and muscarinic type potassium currents that specifically allow OLM cells to exhibit spiking resonance at theta frequencies. Further, we developed a robust Kalman Filtering (KF) method to estimate parameters of the reduced model in real-time. We showed that it may be possible to directly estimate conductance parameters from experiments since this KF method can reliably extract parameter values from model voltage recordings. Overall, our work showcases how the contribution of cellular biophysical current details could be determined and assessed for spiking resonance. As well, our work shows that it may be possible to directly extract these parameters from current clamp voltage recordings.

## 1 INTRODUCTION

It is challenging not only to classify the multitude of different cell types, but also to understand their contributions in brain circuits under normal and pathological states (Zeng and Sanes, 2017; Fishell and Kepecs, 2020). While it is currently possible to record from different cell types *in vivo*, this is technically difficult and laborious to achieve for identified cell types in large numbers. Moreover, a cell’s biophysical characteristics necessarily have to be obtained from *in vitro* studies. Neuronal modelling can bring *in vitro* and *in vivo* studies together by computationally creating artificial *in vivo* states with mathematical models of a given cell type (Destexhe et al., 2003). These models can be viewed as virtual *in vivo* brain circuits. Taking advantage of this approach, we have previously used our computational models to show specialized contributions of interneuron-specific inhibitory cell types in the creation of temporally precise coordination of modulatory pathways (Guet-McCreight and Skinner, 2021). In this way, computational models can help us to gain insight into how specific cell types contribute in different brain states *in vivo*.

Oriens-lacunosum/moleculare (OLM) cells are inhibitory cell types in the hippocampus that function to gate sensory and contextual information in the CA1 (Leão et al., 2012) and support fear memory acquisition (Lovett-Barron et al., 2014). Their firing is phase-locked to the prominent theta rhythms of behaving animals (Katona et al., 2014; Klausberger et al., 2003; Varga et al., 2012). OLM cells have the ability to spike at theta frequencies (Maccaferri and McBain, 1996) and to have a spiking preference to theta frequency sinusoidal inputs (Pike et al., 2000) *in vitro*, although they exhibit little if any subthreshold resonance at theta frequencies (Kispersky et al., 2012; Zemankovics et al., 2010). It is unlikely that OLM cells play a theta pacemaking role since experiments by Kispersky et al. (2012) have shown that OLM cells do not fire preferentially at theta frequencies when injected with artificial synaptic inputs to mimic *in vivo* states. However, if frequency-modulated inputs are presented instead, then there is a theta frequency firing preference. This suggests that OLM cells could contribute to theta rhythms by amplifying theta-modulated activity from presynaptic sources due to their ability to phase-lock with theta-modulated inputs - i.e., they exhibit theta frequency spiking resonance. What biophysical characteristics possessed by OLM cells might be essential to allow them to exhibit spiking resonance at theta frequencies?

A long-known prominent feature of OLM cells is their ‘sag’ which is due to the presence of hyperpolarization-activated cation channels or h-channels (Maccaferri and McBain, 1996). However, in their experimental work, Kispersky et al. (2012) found that theta frequency spiking resonance did not depend on h-channel currents, but it did depend on an after-hyperpolarization (AHP)-like current. OLM cells have a distinct cholinergic fingerprint (Lawrence, 2008; Pancotti and Topolnik, 2022), possessing muscarinic (M-) potassium channel currents that can contribute to AHP behaviours (Lawrence et al., 2006b). Interestingly, muscarinic acetylcholine receptor (mAChR) activation (which inhibits M-channels) of OLM cells has been shown to enhance firing reliablility and precision to theta frequency input (Lawrence et al., 2006a).

Kispersky et al. (2012) generated *in vivo*-like scenarios in their slice preparations using dynamic clamp technology which limited the artificial synaptic inputs to somatic locations. However, synaptic input mostly occurs in dendritic regions. Given this, Sekulić and Skinner (2017) used a developed database of detailed multi-compartment OLM model cells which either had h-channels in their dendrites or not, and created *in vivo*-like scenarios but with artificial synapses that included dendritic regions. They found that OLM cells could be recruited by high or low theta frequency inputs that was dependent on whether h-channels were present in the OLM model cell dendrites or not, respectively (Sekulić and Skinner, 2017). Following this, tightly integrated experimental and modeling work demonstrated that h-channels must necessarily be present on OLM cell dendrites to be able to match experiments (Sekulić et al., 2020). Moving forward, these OLM cell models were used to produce *in vivo*-like (IVL) states, but with consideration of actual presynaptic cell populations (Guet-McCreight and Skinner, 2020). That is, actual synapses were modelled with determined characteristics based on known presynaptic input to OLM cells. Using these IVL states, we showed that spiking resonance at theta frequencies is possible in OLM cells (Guet-McCreight and Skinner, 2021). However, we were limited in the IVL frequencies that our models could express and the detailed multi-compartment nature of the model with thousands of presynaptic inputs prevented a full exploration of what OLM cell biophysical characteristics during IVL states might be critical in bringing about theta frequency spiking resonances.

To tackle this, we here develop a reduced biophysically faithful single compartment OLM cell model and consider this model embedded in a virtual network as represented by IVL states. We use these IVL virtual networks to determine what biophysical details of OLM cells matter for them to exhibit theta frequency spiking resonance. We first ‘validate’ that our reduced model is capable of capturing complex behaviours by showing that it can match results expressed by the full multi-compartment models (Guet-McCreight and Skinner, 2021), and we then examine spiking resonance in IVL states. From an extensive exploration, we are able to show that a combination of h- and M-channels produce the controlling currents for theta frequency spiking resonance. Inspired by Azzalini et al. (2022), we then move on to adjust a robust Kalman Filter (KF) algorithm and use it to estimate parameters of the reduced OLM cell model from membrane potential recordings of the model cells. This indicates that it should be possible to extract parameter values directly from experimental recordings in a real-time fashion. Overall, our work shows that linking *in vitro, in vivo* experimental, computational, and engineering techniques can bring about novel ways to obtain biophysical, cellular understandings of brain circuits.

## 2 METHODS

We used two models in this work. The first model is of our previously published full multi-compartment OLM cell model (Sekulić et al., 2020), but with some minor modifications described below. We refer to this slightly modified model as“FULL”. The second model is a reduced single compartment model that is derived from FULL, and is described in detail below. We refer to it as “SINGLE”.

### 2.1 Multi-compartment OLM cell model

All of the details of our previously developed multi-compartment model can be found in Sekulić et al. (2020). This detailed model was developed in direct and tight correspondence with experimental data and we have referred to it as a “neuron-to-model matched” (NMM) model in our previous work (Guet-McCreight and Skinner, 2021). Here, we use *Cell 1* of the previously developed NMM models. There are nine different currents in this model and we refer to them by their bracketed acronyms: hyperpolarization-activated cation current (Ih), transient sodium (INa), fast delayed rectifier (IKdrf), slow delayed rectifier (IKdrs), A-type potassium (IKa), M-type (IM), T-type calcium (ICaT), L-type calcium (ICaL), calcium-dependent potassium (IKCa).

In examining the multi-compartment model for subsequent reduction, we first simplified the representation of the calcium dynamics to obtain a slightly modified multi-compartment model that we refer to as FULL. The specifics of this modification are provided in SUPPLEMENTARY Material. We note that there is minimal difference (on average less than 1.5 ms difference in spike timing, and less than 0.6% in the resulting voltage and less than 0.12% for any of the biophysical currents in comparing 30, 60, −120 pA current steps) between FULL and the previously published *Cell 1 NMM model*. The set of equations for all the biophysical currents of FULL are provided in SUPPLEMENTARY Material.

### 2.2 Single compartment OLM cell model development

Starting from FULL, we created a single compartment OLM cell model in which biophysical fidelity was maintained as best as possible. In examining all the currents in FULL, we noticed that IKdrs could be removed without much effect. Specifically, we found that there is less than 0.35% difference when comparing voltage peaks (30 and 60 pA current steps) and less than 0.15% for any of the other currents when the IKdrs conductance was set to zero, and less than 0.0015% for voltage or current differences at −120 pA step. Also, we note that in order to be able to capture any of the calcium dynamics in FULL, at least a two-compartment model would be required since calcium channels are only present in the dendrites of FULL - see SUPPLEMENTARY Material for further details.

For practical reasons of: (i) computational efficiency in doing extensive spiking resonance explorations, and (ii) evaluating parameter estimation techniques for direct use with experimental OLM cells, we aimed to have a single compartment mathematical model to use. We thus made the decision to remove consideration of calcium at this stage to enable these practicalities. Therefore, ICaT, ICaL and IKCa were not included in the single compartment model. There are five remaining currents (IKa, IKdrf, IM, Ih, INa) and we proceeded with them to create a single compartment model with biophysical fidelity as follows.

As this *Cell 1 NMM model* was directly matched with experimental data in which both Ih characteristics and passive properties were obtained in the same cell, we created a single compartment model in which these aspects were maintained. Specifically, the surface area, passive properties (leak current, *I_L_*) and Ih, as directly obtained in *Cell 1* (model and experiment) (Sekulić et al., 2020), were minimally changed. The resulting input resistance, time constant and total Ih are comparable to FULL. Further details are provided in SUPPLEMENTARY Material. The conductances of the other four currents, IKa, IKdrf, IM and INa were optimized using BluePyOpt (Van Geit et al., 2016) based on comparison with FULL. Optimization details are provided in SUPPLEMENTARY Material. We refer to this single compartment model with optimized values as SINGLE, and it constitutes our developed single compartment model that is used in this work.

The contribution of the different currents to cell firing is shown in FIGURE 1 for a 60 pA current step for FULL and SINGLE. As easily seen using the currentscape visualization tool (Alonso and Marder, 2019), the relative contribution of the various currents are similar. Although similar, one can observe that there is some compensation for the missing IKCa and calcium currents via IM and IKdrs. These similar balances are apparent at other step currents as shown in SUPPL FIGURE S4. We reasonably claim that SINGLE is a single compartment OLM cell model that maintains biophysical fidelity.

**Figure 1.**
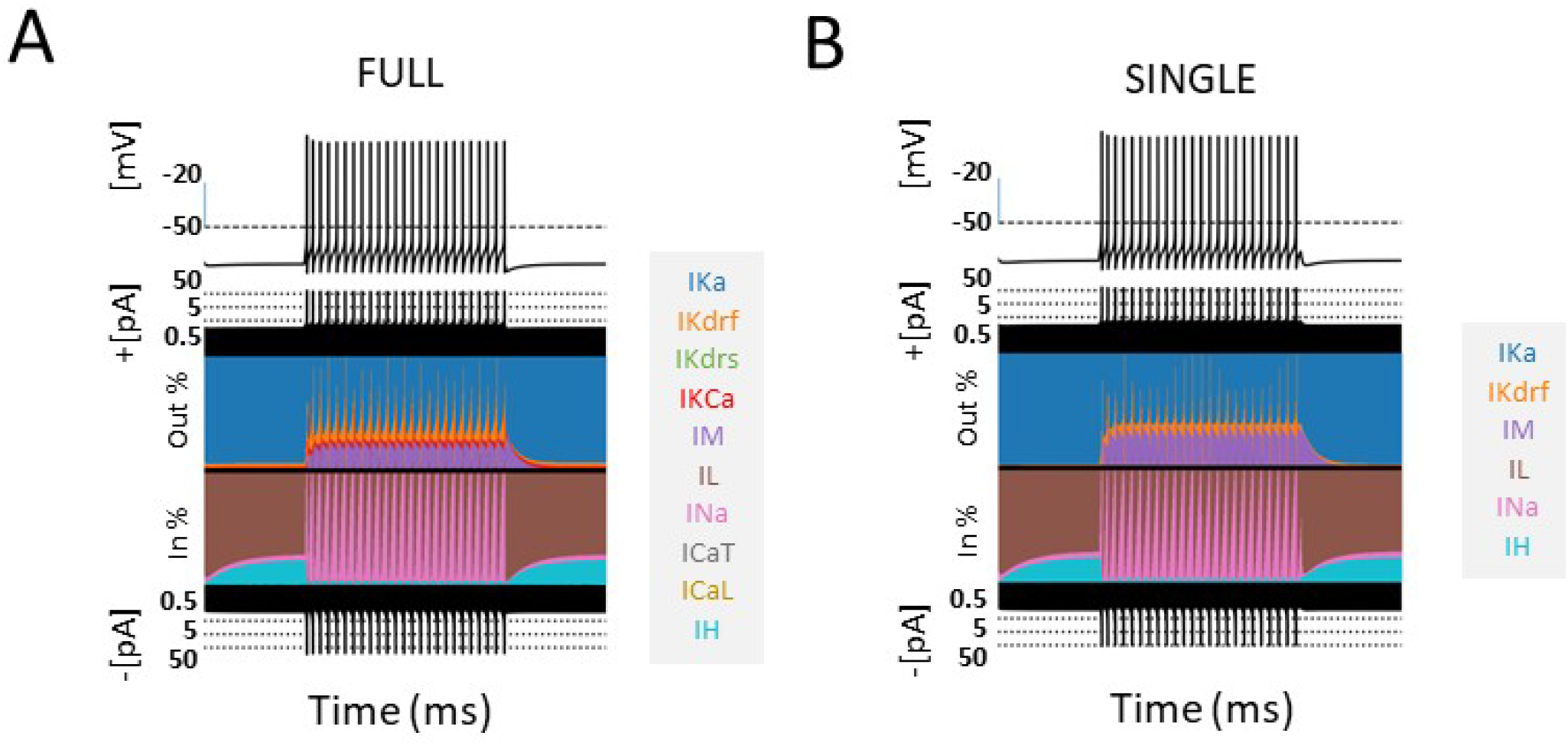
Comparison of models - FULL and SINGLE. Currentscapes (Alonso and Marder, 2019) are shown for models of FULL (somatic location) **(A)** and SINGLE **(B)**. Their firings are similar and corresponding biophysical currents have similar balances. Simulations were conducted for 4000 ms, with a holding current of 4 pA throughout. A step current of 60 pA was injected at 1000 ms for 2000 ms. Comparisons at other step currents are provided in a SUPPLEMENTARY Figure.

### 2.3 SINGLE analyses

#### 2.3.1 Synaptic perturbation model

Previously, Guet-McCreight and Skinner (2021) applied synaptic perturbations to their detailed multi-compartment model, which as noted above is essentially the same as FULL and so we will not distinguish them moving forward. These perturbations were mostly done using 30 (inhibitory or excitatory) synapses randomly distributed across the dendritic tree as a reasonable capturing of experimental observations. Synaptic weights had been optimized based on experiment (Guet-McCreight and Skinner, 2020). Since SINGLE only has one compartment, perturbation was set to be 30 times the individual synaptic weight to maintain an approximately equivalent response in SINGLE relative to FULL. Synaptic equations, implementation and parameter values are identical to *Equation 1* in Guet-McCreight and Skinner (2020) except for the synaptic weight values that are 0.006 (excitatory) and 0.0054 (inhibitory) *μ*S here.

#### 2.3.2 Direct comparisons with FULL

##### 2.3.2.1 In vitro setup and phase response curve (PRC) analyses

###### 2.3.2.1.1 Firing rates

As done for FULL in Guet-McCreight and Skinner (2021), different firing frequencies in *in vitro* states were obtained by injecting constant currents into the cell to produce stable firing rates and inter-spike intervals (ISIs). Firing rates were determined by counting spikes over a 10 second interval.

###### 2.3.2.1.2 PRCs

This analysis was conducted using inhibitory synaptic perturbations with the same setup as in Guet-McCreight and Skinner (2021). That is, we calculated percent phase change in spiking behavior (Δ*ϕ*) from the length of the ISI before the perturbation (T0) and with the perturbation (T1):

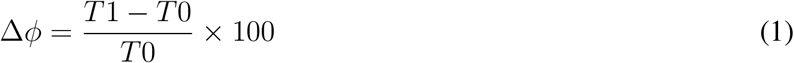

A positive value indicates shortening of the ISI while a negative value indicates a prolonging of the ISI. That is, a phase advance or phase delay respectively. The PRC is generated using perturbations at the full range of phases. To quantify the contribution of the various currents to the PRCs, we also calculated percent change in currents (ΔI) as done in Guet-McCreight and Skinner (2021). For each current, we obtained its maximum amplitude between the 2nd last spike preceding the perturbation (I0), and its maximum amplitude between the perturbation and the second spike following the perturbation (I1). The percent change in maximum amplitude due to perturbation (ΔI) was calculated as:

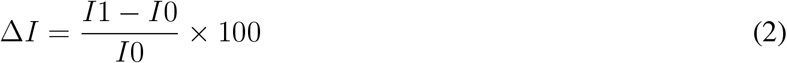

A positive value indicates an increase in maximum current amplitude while a negative value indicates a decrease.

##### 2.3.2.2 Spiking resonance calculations

In Guet-McCreight and Skinner (2021), we computed spiking resonances for *in vitro* and *in vivo*-like (IVL) states. We computed spiking resonances for SINGLE using the same measure as described below.

Each voltage trace obtained was converted to a spike train (1 at spike onset and 0 elsewhere), and the power spectral density (PSD) of each spike train was calculated using the welch function from the Scipy signal module: signal.welch(signal, fs=1/dt, scaling=’density’, nperseg=20000). We defined a measure called the ‘baseline ratio’ to quantify how much a perturbation changes the PSD value from its baseline state. The baseline ratio (*δPSD*) was calculated as:

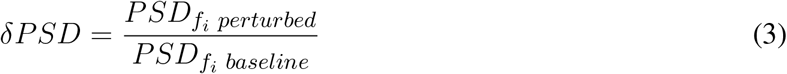

where *f_i_* is the frequency of the input perturbation. That is, 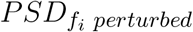 is the spectral density value of the *f_i_* perturbed state spike train, while 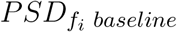 is the spectral density value of the baseline state spike train when there is no perturbation (or a perturbation of ”0”). Thus, *δPSD* measures the effectiveness of an *f_i_* perturbation frequency entraining the cell to fire at that frequency. A value of 1 indicates no effect, while higher values suggest more effective entrainment. The resonant frequency (*f_r_*) is defined as the perturbation frequency that produces the largest *δPSD* value.

###### 2.3.2.2.1 In vitro state

With holding currents between 25.4 pA to 156.8 pA, we generated 50 baseline firing rates from 1 to 36 Hz. For each of the firing rates, we perturbed the voltage using either excitatory or inhibitory synaptic input. As we were doing a direct comparison, we used the same 15 perturbation frequencies as Guet-McCreight and Skinner (2021) that ranged from 0.5 to 30 Hz.

###### 2.3.2.2.2 In vivo-like (IVL) state

A much wider range of baseline firing rates in IVL states can be obtained with SINGLE relative to FULL. For direct comparison with spiking resonances obtained using the multi-compartment model, we restricted the baseline frequencies of SINGLE as such and used the same 15 perturbation frequencies that ranged from 0.5 to 30 Hz.

#### 2.3.3 Expanded explorations with SINGLE

##### 2.3.3.1 In vivo-like (IVL) states with SINGLE

To define an IVL state, we used the same metric as in Guet-McCreight and Skinner (2020) which was previously developed to reflect literature statistics of the OLM cell type in an IVL state. That is:

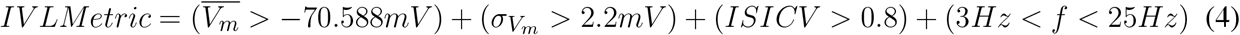

in which the average sub-threshold voltage or membrane potential, 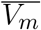, the standard deviation of the sub-threshold membrane potential, 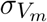, the coefficient of variation for inter-spike intervals, *ISICV*, and the firing rate, *f*, are computed.

In Guet-McCreight and Skinner (2021), several IVL states were obtained with the multi-compartment model by randomizing the synaptic locations on the dendrites. This is of course not possible with SINGLE as it is a single compartment model. Instead, we followed the approach of Destexhe et al. (2001) that implements stochastic synapses via an Ornstein-Uhlenbeck process (Uhlenbeck and Ornstein, 1930). It is formulated as:

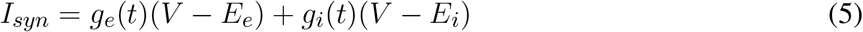

where *g_e_*(*t*) and *g_i_*(*t*) are excitatory and inhibitory conductance values as functions of time. *E_e_* and *E_i_* are excitatory and inhibitory reversal potentials, and *g_e_*(*t*) and *g_i_*(*t*) are given by:

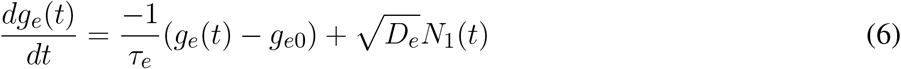

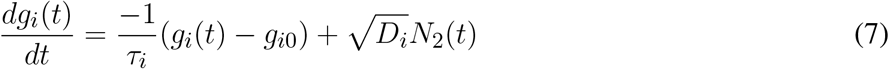

where *τ_e_* and *τ_i_* are decay time constants; *g_e_*_0_ and *g_i_*_0_ are steady state, or average conductances; *N*_1_(*t*) and *N*_2_(*t*) are Gaussian white noise following a standard normal distribution; 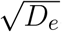and 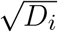 are standard deviations of the noise generated.

###### 2.3.3.1.1 IVL state parameter search

We note that while this stochastic synaptic model would allow several IVL states to be generated with SINGLE, most of its parameters are not directly comparable to IVL states generated with FULL, and as such, we cannot use pre-existing parameter values obtained for IVL states and instead must search for them. We used a brute force approach with nested for loops to search for sets of parameter values for *g_e_*_0_, *g_i_*_0_, 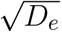 and 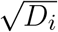 that would generate IVL states. For each set of parameter values, a 10 second long simulation was performed and the resulting voltage trace was evaluated by the IVL metric.

###### 2.3.3.1.2 Constrained search

After experimenting with several parameter range values, we were able to narrow values down to smaller ranges that would mostly output IVL states. The smaller range exploration had *g_e_*_0_ varying from 0.003 to 0.006 with a resolution of 0.0002 μS, *g_i_*_0_ varying from 0.008 to 0.012 μS with resolution of 0.00027 μS, 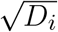 varying from 0 to 0.0016 μS with resolution of 0.000107 μS, and 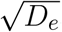 varying from 0 to 0.01 μS with resolution of 0.00067 μS. In addition, we further constrained our IVL states to be comparable with those obtained with FULL. From an analysis of voltage traces of IVL states with FULL from Guet-McCreight and Skinner (2020), we obtained further constraints of: 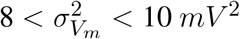 and 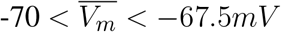. This additional constraint was applied only once to every parameter set to allow output that was close enough to that obtained with FULL.

###### 2.3.3.1.3 Selecting representative IVL states

For each IVL parameter set, fifty 10 second long simulations were run with 50 distinct seeds (the same seeds were used for each set). From these 50 simulations for each IVL parameter set, the minimum and maximum firing rates were recorded, and each set would be clustered as a two-dimensional (2D) system when plotted using its minimum vs maximum firing rate. From this 2D system, 10 IVL parameter sets that span the firing rate space were selected as representative IVL states encompassing the range of firing rate frequencies observed *in vivo* for OLM cells. Since 50 distinct seeds were used, we would have 500 IVL states. For each of these 500 states, we computed firing rate (*f* ’s) and resonant frequencies (*f_r_*’s). *f* ’s were computed as the number of spikes occurring in the 10 second simulation divided by 10, and *f_r_*’s were computed as described above. We removed timepoints occurring at 7 ms on either side of the peak of each spike, and the mean and standard deviation of the remaining (independent) timepoints after spike removal were used to obtain 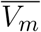 and 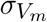. It is possible that some of these 500 states may not fully be IVL states (as defined by the metric) since due to stochasticity the same parameter set might not always generate a (further constrained) IVL state for each random seed used. Based on plots of 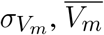 and *ISICV* when using thousands of seeds, the majority of the resulting states remain as IVL states (see SUPPL FIGURE S7).

##### 2.3.3.2 Spike Triggering Average (STA) analyses

To extract biophysical contributions of resonant frequencies in IVL states, we used STA analyses (Ito, 2015; Schwartz et al., 2006). Oftentimes, a STA analysis (or reverse correlation) is applied using mean input currents that precede spikes, and in a comparative fashion with different cells types in experiments to gain insight (e.g., see Moradi Chameh et al. (2021).) Here, because we are using mathematical models, we can consider relative contributions of the underlying active and passive currents preceding spikes in a comparative fashion. This is in an analogous vein to when we considered how the different currents responded with phase-dependent perturbations in addition to the ‘standard’ PRC obtained when one considers how much the subsequent spike is advanced or delayed given phase-dependent perturbations.

For spiking resonances, we used 27 different perturbation frequencies. For direct comparisons of spiking resonances in IVL states between FULL and SINGLE we used the same set of perturbation frequencies (15) as used in Guet-McCreight and Skinner (2021). However, with a much wider range of baseline firing rates in IVL states with SINGLE, we could obtain a larger set of spike resonant frequencies using the larger number of different perturbation frequencies.

###### 2.3.3.2.1 Generating STA plots

We define a set of frequencies *F* = {0.5} ∪ {1 ≤ *i* ≤ 25, *i* ∈ ℤ} ∪ {30} Hz. To obtain a spiking resonant frequency, *f_r_*, in *F*, we ran 10 second long simulations using the 10 representative parameter sets to find IVL states where the model was resonant to that frequency. For each representative set, each of the 10 second long simulations were produced with a different seed. Histogram distributions of the different *f_r_*’s naturally depend on how many random seeds one uses. [[Refer to OSF FIGURE that shows these distributions for 500 and 5000 seeds.]] With enough seeds, we were able to obtain *f_r_* for all of the different frequencies in *F* for each representative parameter set.

We generated STA plots in which we considered up to 200 ms preceding a spike. To do this, we found as many 200+ ms long ISIs as possible within each 10 second long simulation. We used new seeds to run our simulations until there were 50 (200+ ms ISIs) for each representative set of *f_r_*’s for all of the 27 frequencies in F. The actual number of seeds required ranged from 40,000 to 120,000 differing across the 10 representative sets. For the 200 ms ISIs we obtained, we preserved sections from 195 ms to 10 ms before each spike so as to remove the influence from the last and the next spike on the underlying currents. As there is ‘curving up’ before a spike in the STA plots (see SUPPL FIGURE S8) that could drastically affect the slope analysis (see below), we chose a timepoint range of 195 ms to 25 ms before a spike. 195 ms is justified to be approximately the furthest back one can push to avoid the previous spike, and 25 ms is approximately the furthest forward one can push to not include the curving portion before slope computation. For computational ease, we used this same range in doing both the slope and spread analyses described below. With these ISIs, we generated STA plots to consider the contribution of the different biophysical currents.

The biophysical currents in the STA plots were produced from a *f_r_* and representative set (*set*#) perspective. That is, for each current type *I_x_*, there were 270 different average (of 50) *I_x_*’s for each timepoint from −195 to −25 ms, each corresponding to a permutation of *f_r_* value (27) and *set*# (10). The biophysical current value itself represents the proportion of the total (absolute) current at the given timepoint where it is assigned as negative (inward current) or positive (outward current) accordingly.

Referring to these currents as 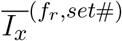, the average current (of 50), *f_r_* ∈ *F*, and 0 ≤ *set*# ≤ 9, we took the absolute values of the averages, and normalized them across the representative sets to keep the values of each current between 0 and 1. We refer to normalized 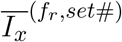 as 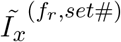. That is, we now have:

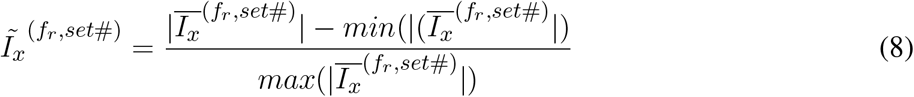

In Figure 7A, a selection of 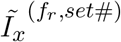 are shown.

###### 2.3.3.2.2 Slope Analysis

For each timepoint of 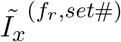, we calculated the mean across the 10 representative sets. These means can be seen in FIGURE 7B with the standard deviations.

For each *f_r_*, we did a least squares linear fit line of the mean, and obtained the slope of the linear fit line. We also considered a combined slope of IM and Ih where the two currents were first summed before obtaining the mean and subsequent slope from the linear fit.

###### 2.3.3.2.3 Spread Analysis

For each timepoint of 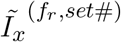, we calculated the standard deviation across the representative sets. We defined the function 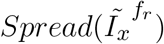 as twice the standard deviation summed over all the timepoints from 195 to 25 ms before a spike. In FIGURE 7B, the precursor to 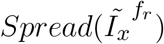, the mean and standard deviations of 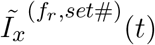 are shown for a selected set of *f_r_*’s.

We also considered a combined spread of two currents IM and Ih, 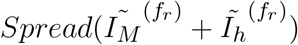, where the two currents were first summed, and then twice the standard deviation of the resulting summation was summed over all the timepoints. Note that with given current type(s), the spread is a function of *f_r_* only as we sum over the timepoints.

### 2.4 Direct parameter estimation

#### 2.4.1 Robust Adaptive Unscented Kalman Filter (RAUKF)

The mathematical model structure of SINGLE may also be used in data assimilation methods to estimate parameters based on an ‘observation’. Application of the RAUKF method (Azzalini et al., 2022) here was developed based on the reduced OLM model cell, and the model parameters were estimated from simulated noisy observations of SINGLE or FULL. The RAUKF method was adapted so that the conductance based model used for state prediction was the same as that used in SINGLE (See details in SUPPLEMENTARY Material). The general formulation of the Kalman Filter (KF) is:

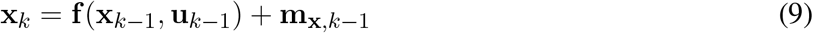

Where the state at *k*, **x**_*k*_ depends upon the previous state **x**_*k*−1_ and some control input **u**_*k*−1_, i.e. injected current, and 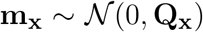. The function **f**is the system’s state model, which in this case are the set of equations governing model SINGLE. In order to estimate parameters of the system, additional trivial dynamics were added to the system for each given parameter ***θ*** via a random walk:

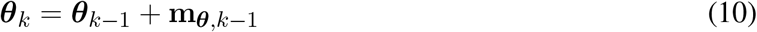

With 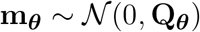. Combining **x** and ***θ*** we may produce a new state **X** with the dynamics represented by **F** and noise **M**, where **I** is the identity function:

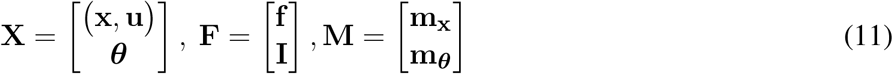

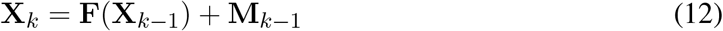

The observation model used to characterize noisy membrane voltage measurements is described by

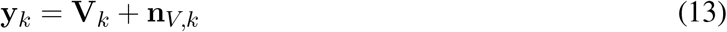

where *n_V,k_* denotes measurement noise (in the general case, 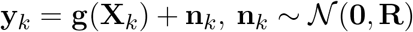). Here, the direct observation of the membrane voltage *y_k_* mimics experimental recording techniques (e.g., current-clamp methods). With only a subset of **X**_*k*_ being measurable, the method presented in this study allows hidden states to be estimated.

The system’s dynamics model **F**, an initial estimate of the state **X**_0_, as well as the control input **u** were used to maximize the probability of an observation **y** afforded by the noise profiles of each.

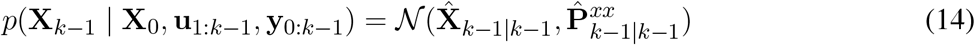

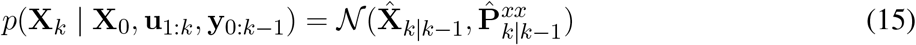

To estimate the next state (15) a set of points (sigmapoints) were sampled according to (14), described in more detail in Azzalini et al. (2022). This process also updates the current covariance **P** of the process.

Since the precise noise profile **Q** of the state vector **X** is not well known for a given neuronal model, and its respective discrepancies to the observation, we employed an adaptive method used in previous work in order to update **Q** as well as the estimate of the observation noise **R** over time (Azzalini et al., 2022). See SUPPLEMENTARY Material for further details.

The adaptation of **Q** and **R** can be tuned by user defined variables *a*, *b*, *λ*_0_, and *δ*_0_, Where *λ*_0_ and *δ*_0_ are the minimum weights used when updating **Q** and **R**, the percent shift from the current to the new estimated value. To ease computational load these updates to **Q** and **R** were not applied after every sample, instead they were applied when a fault was detected (see SUPPLEMENTARY Material). The values of *λ*_0_ and *δ*_0_ were both set to a value of 0.2. The variables *a* and *b* are weights of how much **Q** and **R** will be updated proportional to the magnitude of fault detected. The values of *a* and *b* were set according to explorations done in previous work when RAUKF was applied to a conductance based model (Azzalini et al., 2022). *a* >> *b* was found to be effective and here we use *a* = 10 and *b* = 1.

#### 2.4.2 Application of RAUKF to OLM cell models

When applying the RAUKF to an observation of either SINGLE or FULL, ***θ*** was set to estimate the same conductance values that were optimized in the reduced model SINGLE. That is,

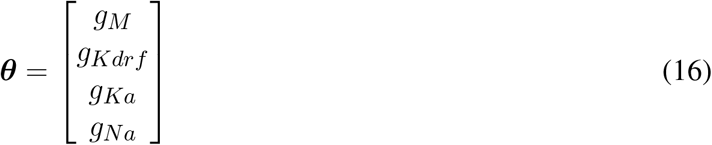

To generate an observation, an input train of step currents was injected into either SINGLE or FULL models. The input train consisted of 500 ms of zero current, then a step to 30 pA, 60 pA, 90 pA, 0 pA, −30 pA, −60 pA, −90 pA, repeating 4 times. The current was also mixed with a Gaussian white noise 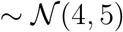. The mean of the noisy stimulation was chosen to match the characterized bias current in FULL and SINGLE. This input train of multiple positive and negative pulses is designed to be able to expose a full dynamic range of spiking, resting and subthreshold activities due to the underlying biophysical currents, and can be used in real-time to estimate parameter values.

The initialization of **P** and **Q** was derived from the order of magnitude ***θ***, while **x** was made constant to 1 × 10^−4^ and 1 × 10^−8^ respectively. For the **P** of ***θ*** the value was set to 2× the magnitude of the value of ***θ*** used in SINGLE, e.g. 340 → 1 × 10^2^ → 1 × 10^4^, and 4× the order of magnitude was used to initialize **Q**. Two initial states were used for ***θ***_0_, one where ***θ***_0_ = ***θ***_*SINGLE*_, known as the optimized initial estimate, and another where 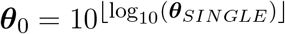, known as the poor initial estimate.

### 2.5 Computing

The extensive SINGLE simulations with spiking resonance and STA analyses were done using the Neuroscience Gateway (NSG) for high-performance computing (Sivagnanam et al., 2013). Code pertaining to FULL, SINGLE and analyses is available at: https://github.com/FKSkinnerLab/Reduced_OLM_example_code. Code pertaining to RAUKF and the respective figures is available at: https://github.com/nsbspl/RAUKF_OLM

## 3 RESULTS

### 3.1 Reduced model can capture complex outputs obtained with full, detailed multi-compartment model

From our detailed multi-compartment OLM cell model (FULL), we derived a reduced, single compartment model (SINGLE) as described in the Methods. While SINGLE can reproduce features of FULL (e.g., see FIGURE 1), it is not obvious that SINGLE would produce similar results to those obtained using the full, detailed multi-compartment model regarding more complex phenomena such as spiking resonance (Guet-McCreight and Skinner, 2021).

Considering this, we used SINGLE to examine previously explored phenomena so as to directly compare with observations obtained using the detailed multi-compartment model. In FIGURE 2, we show that SINGLE and FULL have similar frequency-current (f-I) curves, with the required injected current to obtain about a 1 Hz firing frequency differing by about 4 pA. As described in the Methods, the detailed multi-compartment model used in Guet-McCreight and Skinner (2021) is essentially the same as FULL. Thus, moving forward we will not specifically distinguish these multi-compartment models in comparing our results.

**Figure 2.**
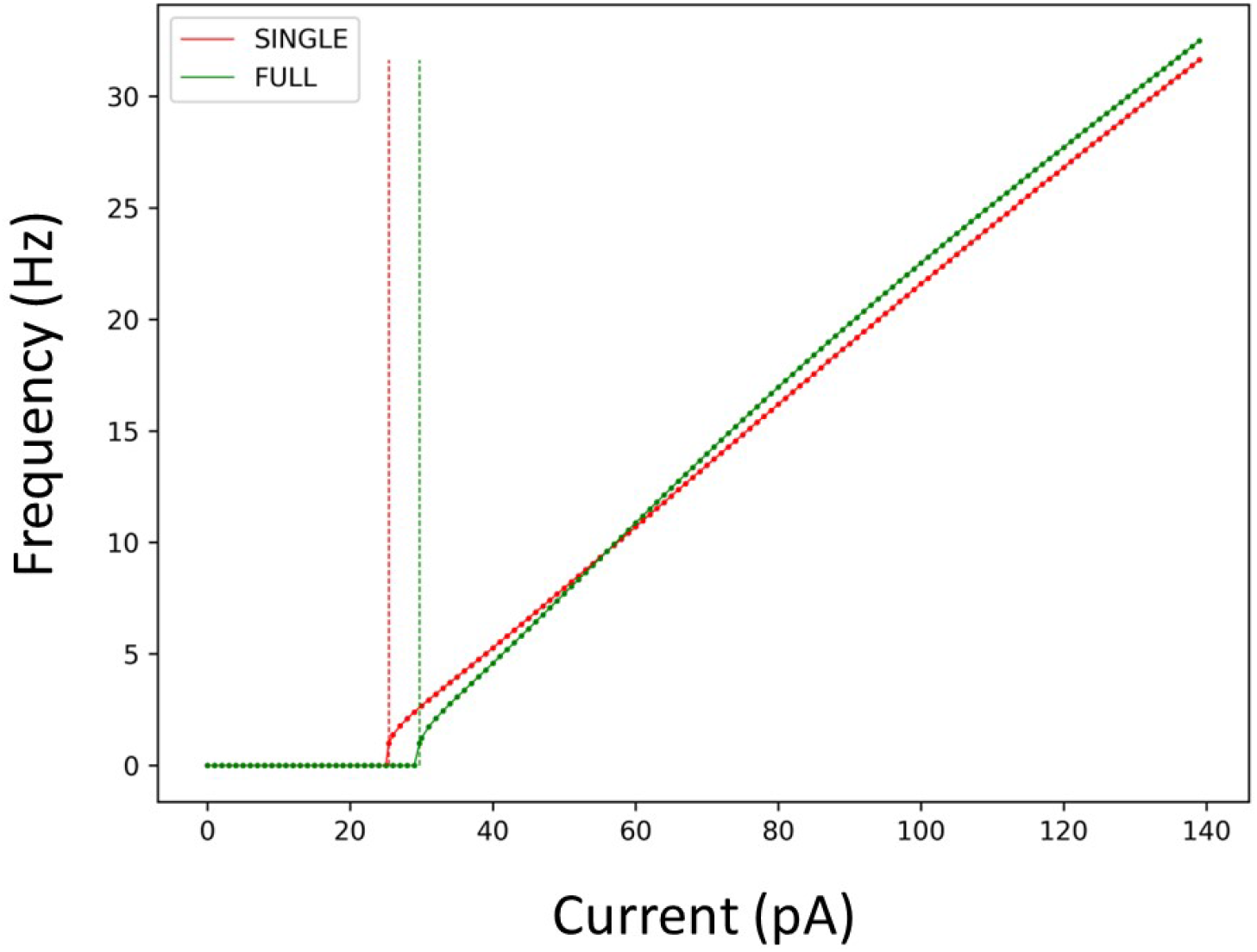
Firing rates of SINGLE and FULL. The red and green dots and lines refer to the frequency vs current (f-I) curves for SINGLE and FULL respectively. Correspondingly, the red and green dashed lines signify the amount of injected current needed to obtain a firing rate of 1 Hz in SINGLE and FULL. 25.5 pA for SINGLE and 29.7 pA for FULL.

We examined phase response curves (PRCs) and the phase-dependent contribution of the different biophysical currents at four different frequencies when inhibitory perturbations are applied, as previously done in Guet-McCreight and Skinner (2021). We show the results in FIGURE 3A-C for the PRCs, Ih and IM which illustrates that similar results are obtained. That is, inhibitory perturbations mostly delay spiking and drive up the responses of Ih and IM. Further, the percent change in maximum Ih increases in magnitude as the firing rate increases, and its peak initially shifts rightward followed by a gradual leftward shift. For IM, the magnitude increases and the negative peak shifts leftward as the firing rate increases. Phase-dependent changes for all the other currents are shown in OSF FIGURES OSF.

**Figure 3.**
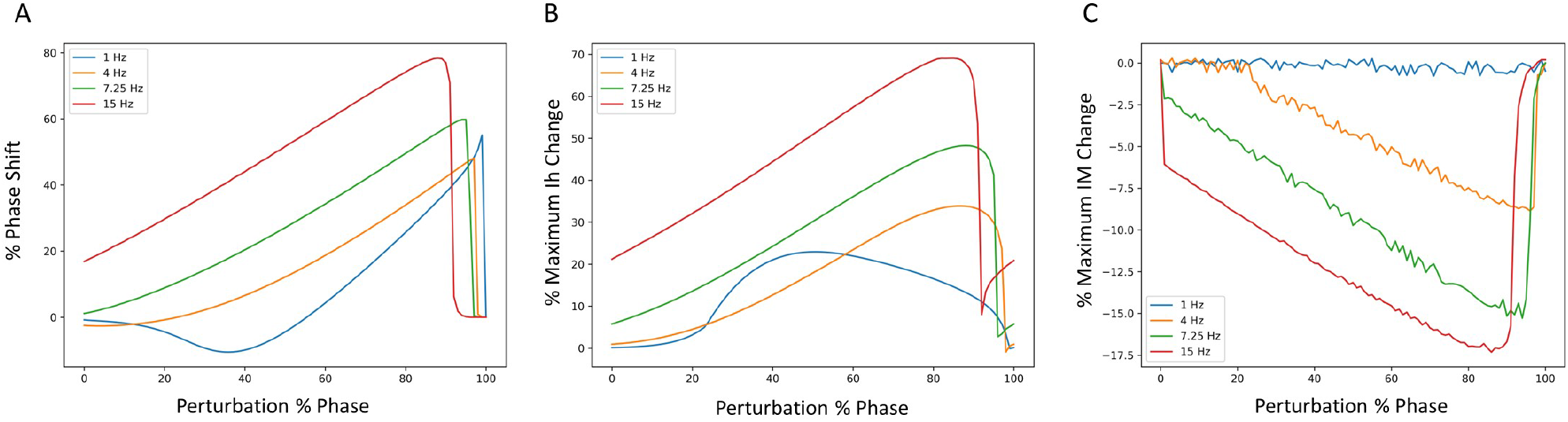
Phase response curves (PRCs) using different firing frequencies of SINGLE. **(A)** shows PRCs at 4 different frequencies and the corresponding phase-dependent changes of IM **(B)** and Ih **(C)** when inhibitory perturbations are applied to SINGLE firing at 1, 4, 7.25 and 15 Hz, due to injected currents of 25.5, 35.1, 47.4, 75.7 pA respectively. Responses are similar to those obtained by (Guet-McCreight and Skinner, 2021).

We also undertook a full examination of spiking resonance in an *in vitro*-like scenario, using the same range of firing frequencies as used in Guet-McCreight and Skinner (2021). A side-by-side comparison of spiking resonance results using SINGLE and the previously published data using a detailed multi-compartment model is shown in FIGURE 4. Again, the results are similar. That is, with inhibitory perturbations, the resonant frequency of both models increases as the baseline firing rate increases (FIGURE 4A). However, with excitatory perturbations, the resonant frequencies mostly lie within the theta frequency range regardless of the baseline firing rate (FIGURE 4B).

**Figure 4.**
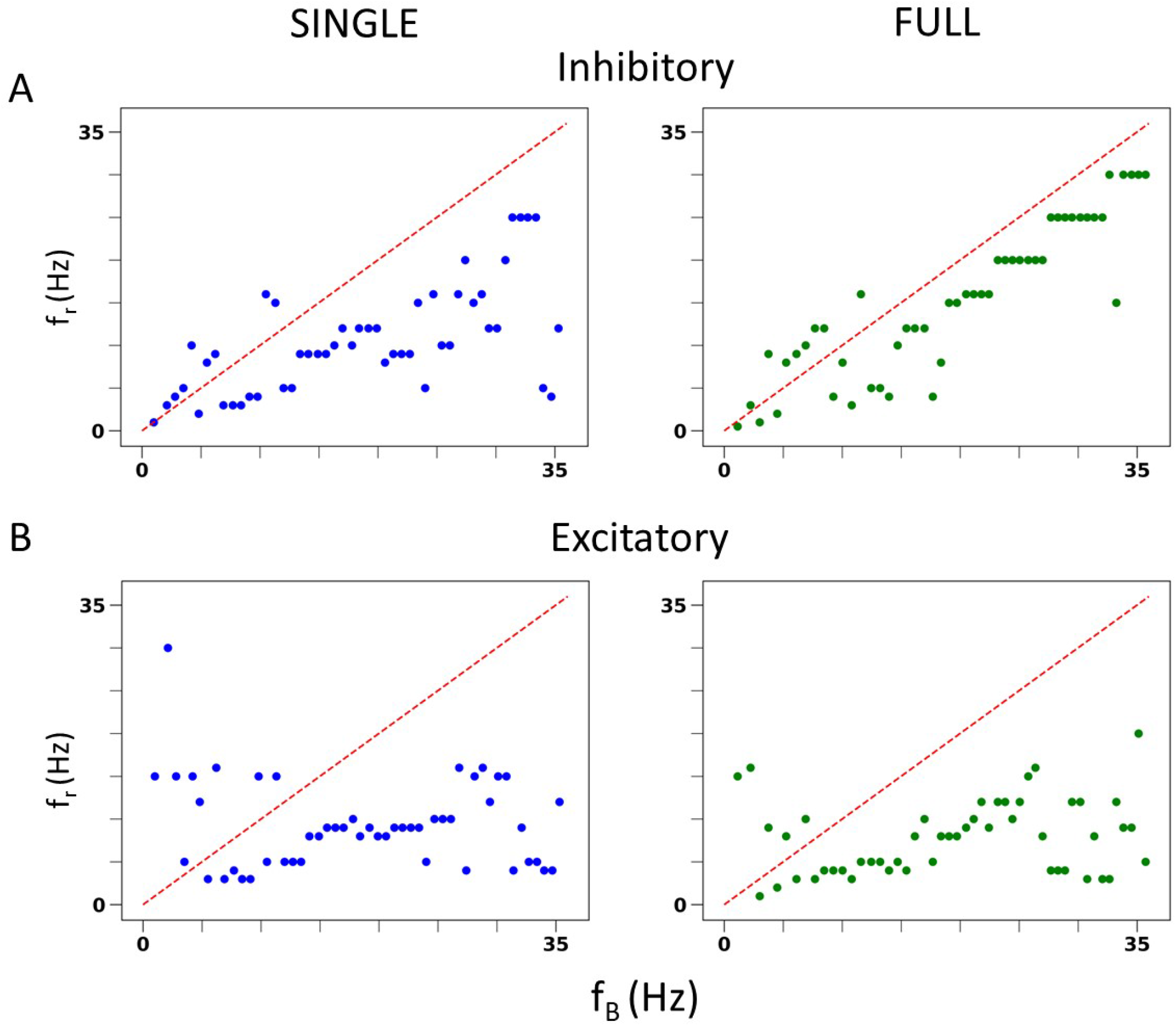
*In vitro* spiking resonance pattern in SINGLE is comparable to FULL. **(A)** Resonant frequency (*f_r_*) of inhibitory perturbation at different baseline firing rates (*f_B_*). Blue is from single compartment model, green is from full model. **(B)** Resonant frequency (*f_r_*) of excitatory perturbation at different baseline firing rates (*f_B_*). Blue is from single compartment model, green is from full model.

As described in the Methods, we can generate *in vivo*-like (IVL) states in our reduced model. Restricting the baseline frequencies to those used in Guet-McCreight and Skinner (2021), similar spiking resonance results in IVL scenarios are also obtained as shown in SUPPL FIGURE S6. That is, excitatory perturbations mostly do not evoke spiking resonance at theta frequencies whereas inhibitory perturbations do. Thus our reduced model, SINGLE, is not diminished in being able to reproduce complex results found in FULL.

### 3.2 Expanded virtual network explorations with reduced model

#### 3.2.1 A wide range of *in vivo*-like (IVL) states is possible with SINGLE

*In vivo* recordings of OLM cells exhibit a range of baseline firing frequencies that span ≈ 3 – 25 Hz (Katona et al., 2014; Varga et al., 2012) (depending on behavioural state of movement, sleep etc.). However, our previous spiking resonance explorations in IVL states using full, detailed multi-compartment models were technically constrained to a small range of baseline firing frequencies, specifically 12 16 Hz (Guet-McCreight and Skinner, 2021), due to how different IVL states could be generated. As such, we were limited in being able to extract biophysical underpinnings of different spike frequency resonances that include theta frequencies. Here, with SINGLE, our reduced model that maintains biophysical balances for the 5 different currents it contains (see FIGURE 1), we are able to obtain a wide range of baseline firing rates and thus can greatly expand our explorations in IVL states. This puts us in a position to determine what the biophysical determinants underlying theta frequency spiking resonance might be.

We obtained a wide range of baseline firing rates with SINGLE by altering the parameters for the ‘noisy’ excitatory and inhibitory currents it receives (see Methods). From our parameter search, 414 different sets of parameters that can generate valid IVL states were found. Within these sets, the excitatory conductance and variance were generally smaller than their inhibitory counterparts, suggesting that inputs received by OLM cells *in vivo* may be inhibitory dominant. From these 414 sets, we chose 10 that could span the full range of *in vivo* firing frequencies of OLM cells, and refer to them as 10 representative parameter sets. Each representative set has a minimum and maximum baseline firing rate as given in TABLE 1 along with its parameter set values.

**Table 1.** Representative Parameter Set Characteristics.

| Set # | Min $f$ | Max $f$ | $g_{e0}$ | $g_{i0}$ | $\sqrt{D_e}$ | $\sqrt{D_i}$ |
| --- | --- | --- | --- | --- | --- | --- |
| 0 | 3.1 | 4.6 | 0.00321429 | 0.01085714 | 0.00068571 | 0.00214286 |
| 1 | 4.6 | 6.8 | 0.00342857 | 0.01028571 | 0.00057143 | 0.00214286 |
| 2 | 6.5 | 9.3 | 0.00342857 | 0.01114286 | 0 | 0.00428571 |
| 3 | 8.3 | 10.9 | 0.00385714 | 0.01171429 | 0 | 0.00428571 |
| 4 | 10.4 | 13 | 0.00428571 | 0.01171429 | 0.00045714 | 0.00357143 |
| 5 | 11.8 | 15.2 | 0.00385714 | 0.01142857 | 0.00011429 | 0.00571429 |
| 6 | 13.3 | 16.8 | 0.00428571 | 0.01114286 | 0 | 0.00428571 |
| 7 | 15.2 | 18.6 | 0.00407143 | 0.01 | 0 | 0.00428571 |
| 8 | 16.7 | 19.9 | 0.00364286 | 0.00828571 | 0 | 0.00428571 |
| 9 | 18.5 | 22.6 | 0.0045 | 0.01085714 | 0.00011429 | 0.00571429 |

For each representative set, we generated fifty 10 second long IVL states (via 50 random seeds) to obtain 500 IVL states. In FIGURE 5, we plot the minimum and maximum firing rates of these 500 IVL states. To illustrate what the OLM cell firings in IVL states look like, one example of an IVL state from each of the 10 chosen representative sets is also shown in FIGURE 5.

**Figure 5.**
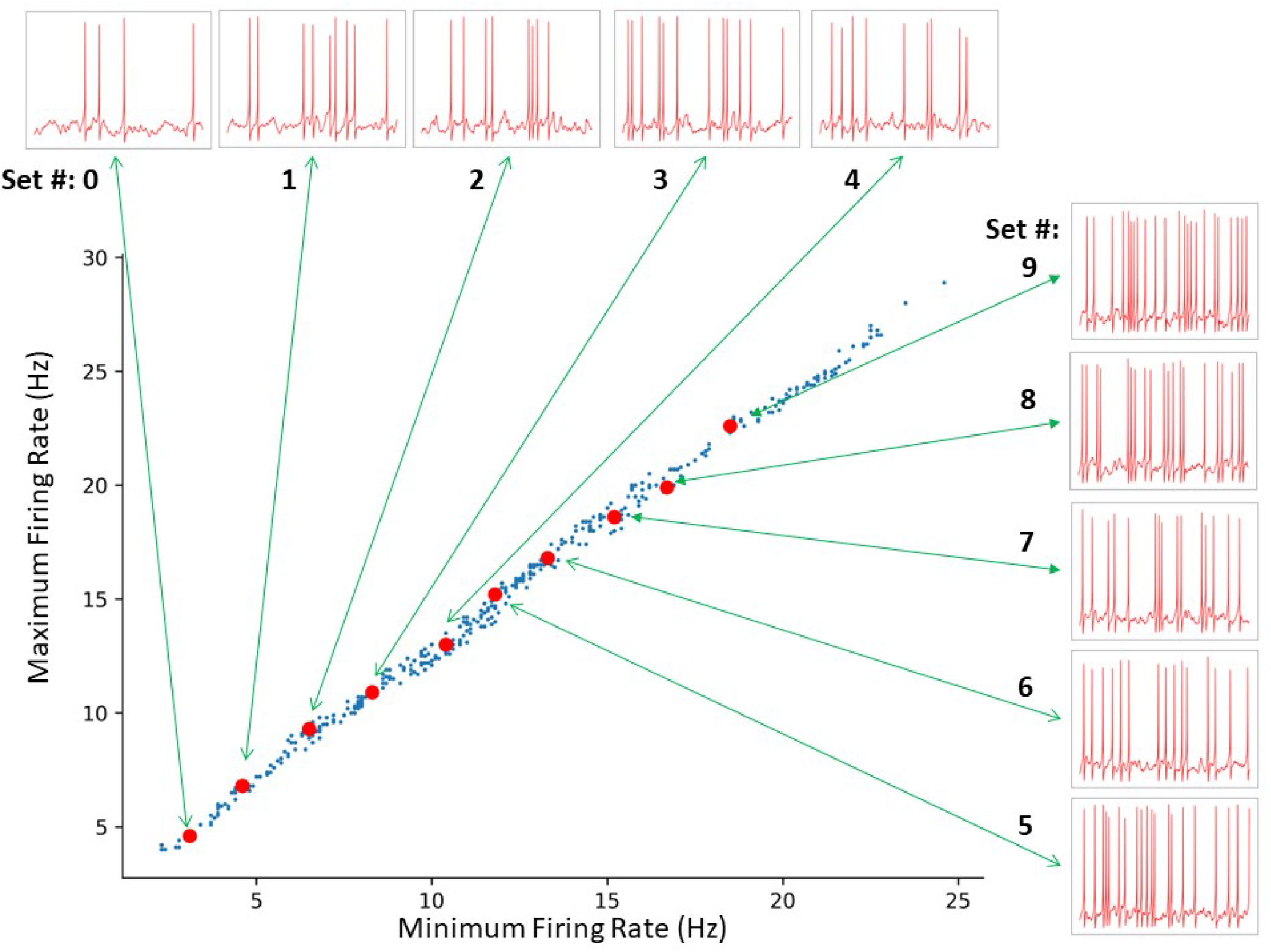
SINGLE yields a wide range of *in vivo*-like (IVL) firing frequencies as exists in experiment. A total of 414 sets of parameters were identified by a constrained search. The red dots indicate the 10 representative sets selected. From bottom left to top right: set 0 to 9, respectively. The green arrows connect each representative set to a one second voltage trace example produced using the respective representative set.

We can generate as many IVL states as we desire simply by using >50 random seeds. This was also the case in our previous work using a full multi-compartment model, but the randomness was in regard to synapse placement and presynaptic spike times from specified presynaptic cell types, resulting in a limited baseline firing frequency range (Guet-McCreight and Skinner, 2021). Here, the randomness is less restrictive because of the single compartment model nature of SINGLE and is able to exhibit the needed full range of baseline firing frequencies.

#### 3.2.2 spiking resonances with SINGLE

We are not limited in how many random seeds we can use. We already know that we can capture the full extent of *in vivo* firing ranges, and we aimed to generate a large data set of spike frequency resonances. With a large data set, we anticipated that we could determine whether there are any particular biophysical determinants underlying *theta frequency* spiking resonance specifically. We generated 50,000 IVL states from the 10 representative sets using 5,000 random seeds. In FIGURE 6A, we show the computed spiking resonance (*f_r_*) as a function of the baseline firing frequency (*f_B_*) when we used inhibitory perturbations. The analogous plot for excitatory perturbations is shown in FIGURE 6B. To ensure that the individual dots are visible, we only used the first 50 seeds (500 IVL states) in creating the plots shown in FIGURE 6. As can be seen, *f_B_* spans the range, unlike the case with FULL (see SUPPL FIGURE S6). In SUPPL FIGURE S7, we plot the mean 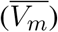 and variance 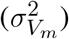 of the sub-threshold membrane potential, and the *ISICV* to show that these aspects of IVL states are not determining factors of the resulting *f_r_*’s. That is, for any given *f_r_*, there are a range of 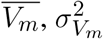 and *ISICV* values. Equivalently, resonant frequencies are distributed across the ranges of 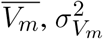 and *ISICV* values.

**Figure 6.**
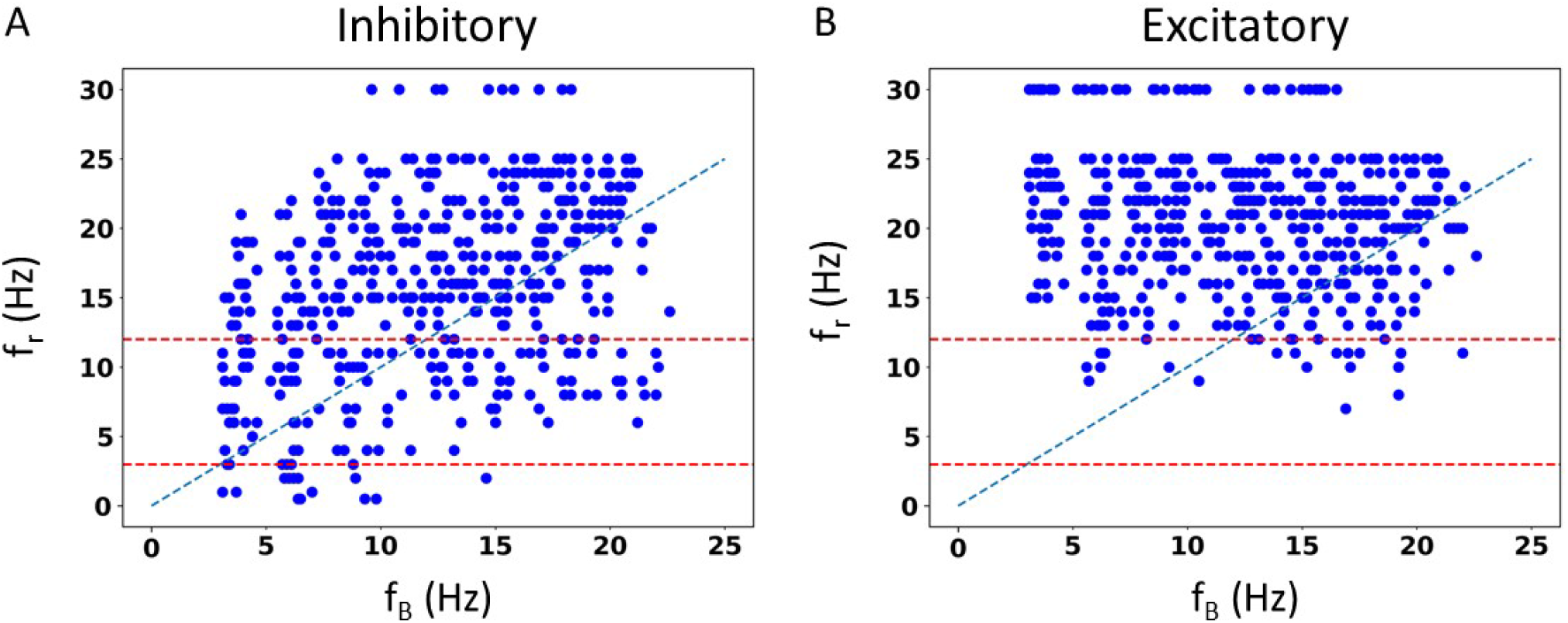
Spike resonances using SINGLE. 500 IVL states generated from the 10 representative sets are used to plot resonant frequencies (*f_r_*’s) as a function of (average) baseline firing rates (*f_B_*’s). Plots are shown for inhibitory perturbations **(A)** or excitatory perturbations **(B)**. Diagonal line is where *f_r_* = *f_B_*. Red dashed lines delineate a theta frequency (3-12 Hz) range for the *f_r_*’s.

Not surprisingly, one can see some dependence of *f_r_* on *f_B_*. That is, with a higher frequency *f_B_*, there is an increase in the density of higher frequency *f_r_*’s. This can be seen in FIGURE 6, as well as in the density of colored dots representing *f_B_* in SUPPL FIGURE S7. That is, IVL states with a higher *f_B_* are more likely to have higher *f_r_*, even though due to stochasticity of the IVL states, it is possible for some higher *f_B_* IVL states to be resonant at low frequencies. However, this observation holds true on average. In essence, we have *f_r_*’s at many different frequencies.

Using FULL, we had previously noted that inhibitory perturbations, and not excitatory perturbations, generated *f_r_*’s that include theta frequencies during IVL states (Guet-McCreight and Skinner, 2021). Here, with SINGLE, this observation can be more clearly seen, as shown in FIGURE 6 where dashed red lines to delineate the theta frequency range (3-12 Hz) are drawn. With our greatly extended dataset, we can easily see that the full theta frequency range of *f_r_* with inhibitory perturbations is exhibited. However, with excitatory perturbations, this is not the case - only minimally and at the upper end of theta frequency range are *f_r_*’s obtained. We note that the limited baseline frequencies available with FULL prevented us from being able to fully observe this.

#### 3.2.3 STA analyses

As we would like to understand what underlies the ability of OLM cells to exhibit spiking resonance at theta frequencies, we now focus on just the inhibitory perturbations as it is inhibitory, but not excitatory, that encompass the full range of theta frequency spiking resonances (see FIGURE 6).

To consider this, we undertake a spike-triggered average (STA) analysis from the perspective of all of the underlying biophysical currents. We examined each current type’s contribution to spiking in IVL states for all of the different *f_r_*’s (27 in total from 0.5 to 30 Hz). In FIGURE 7A, we show STA plots for six different *f_r_*’s from these 27. For each *f_r_*, we show normalized values in the STA plots for the five different biophysical currents (and leak) for each of the ten representative sets (see Methods for details).

**Figure 7.**
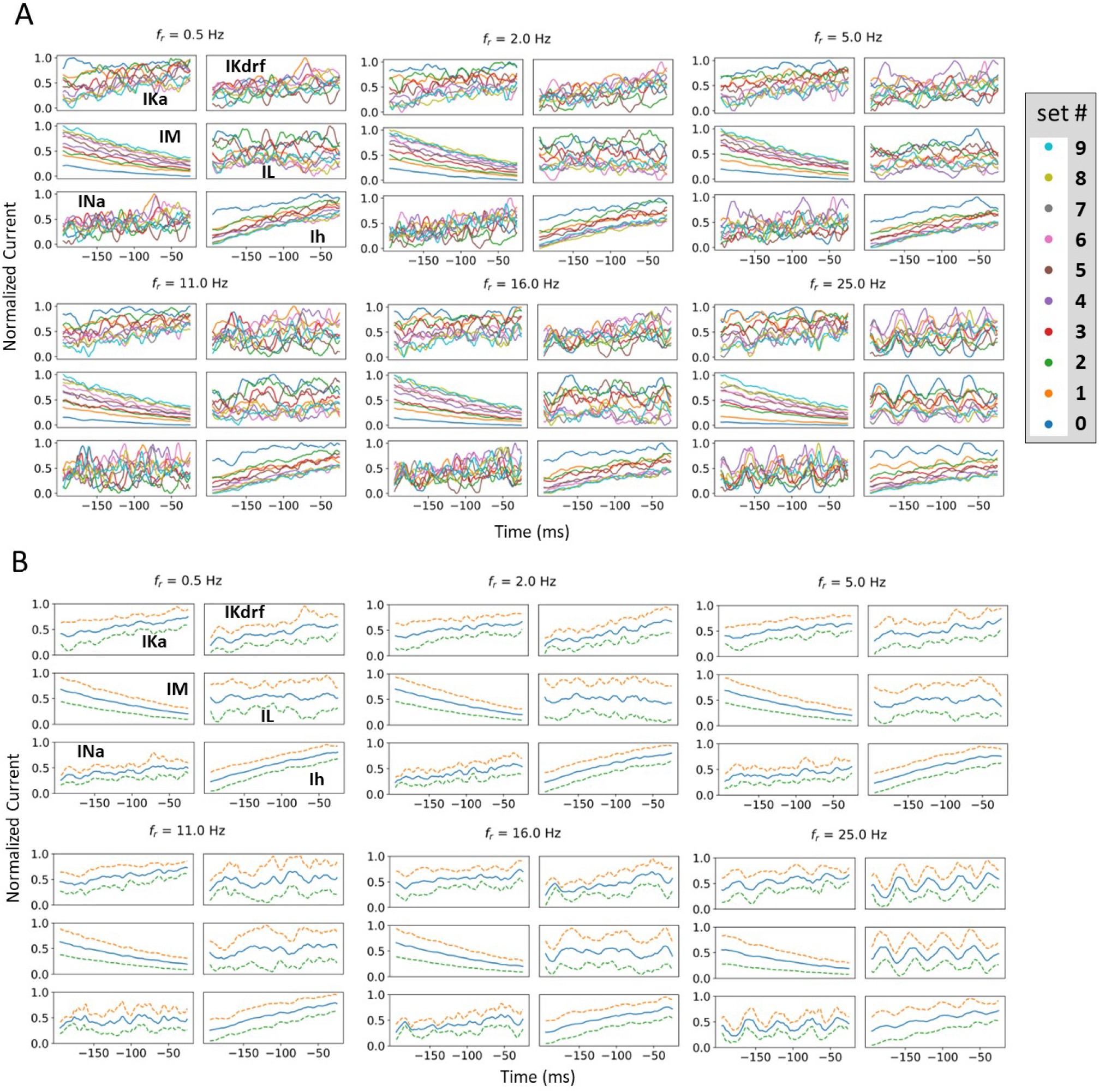
Examples of normalized STA and spread analysis. **(A)** Examples of normalized STA plots, with different colors represent different representative sets. **(B)** Blue lines in the middle of the plots are the means of normalized currents from (A). The dashed lines are ±1 standard deviation of the normalized currents from the mean. Spread is defined as the mean difference between the dashed lines.

From STA plots using un-normalized values, the proportions of currents are directly comparable and it is clear, for example, that the relative proportion of Ih (to the total current) is much smaller than that of IM. From a thorough examination of STA plots of the un-normalized current values (proportion of the total current), we find that the larger the *f_r_*, the more negative is the Ih and the less positive is the IM. In addition, STA plots with higher *f_r_* frequencies show obvious fluctuations. These fluctuations would seem to be due to the presence of more perturbations, with the peaks and valleys of the fluctuations matching the timing of the perturbations. However, IM appears the least sensitive to the perturbations, with negligible fluctuations (see FIGURE 7A). Of particular note is that STA plots of IM and Ih show minimum overlap between the different representative sets relative to the other currents. This is more obviously the case for IM whose overlap is visually separable (see FIGURE 7A.) In general, overlap is likely the result of stochasticity in the OU process which makes the currents variable and intersecting with each other. The minimal overlap in IM and Ih indicates an insensitivity to local changes, which could be attributed to them having slower time constants relative to the others. This is especially true for IM which has the slowest time constant (see equations and plots for the biophysical currents in SUPPLEMENTARY Material). This would also explain why it does not show fluctuations for larger *f_r_* whereas Ih does. A general interpretation of this minimal overlap is that regardless of what the *in vivo* firing rate of the OLM cell might be, IM and Ih can ‘tailor’ their response accordingly. That is, they can perceive “global changes” and so the different firing rates are in the ‘history’ of IM and Ih more than the other current types with faster time constants. Therefore, we now reasonably assume that if there is going to be any frequency specificity in the OLM cell’s spiking resonance, Ih and IM would be the currents that could potentially control this. We now focus on Ih and IM to determine whether there are any particularities associated with theta frequency spiking resonance. To do this, we use an analysis that involves slopes and spreads. See Methods for computation details.

Let us first consider our slope analyses. We found that as *f_r_* increased, the slope of IM’s STA plot became less negative, and the slope of Ih’s became less positive. Linear fits are statistically significant. In considering a combination of IM and Ih slopes, we again obtained a statistically significant linear fit, but now the slope crossed zero within the theta frequency range. This is all shown in FIGURE 8A where the 3-12 Hz theta frequency range is delineated by dashed green lines.

**Figure 8.**
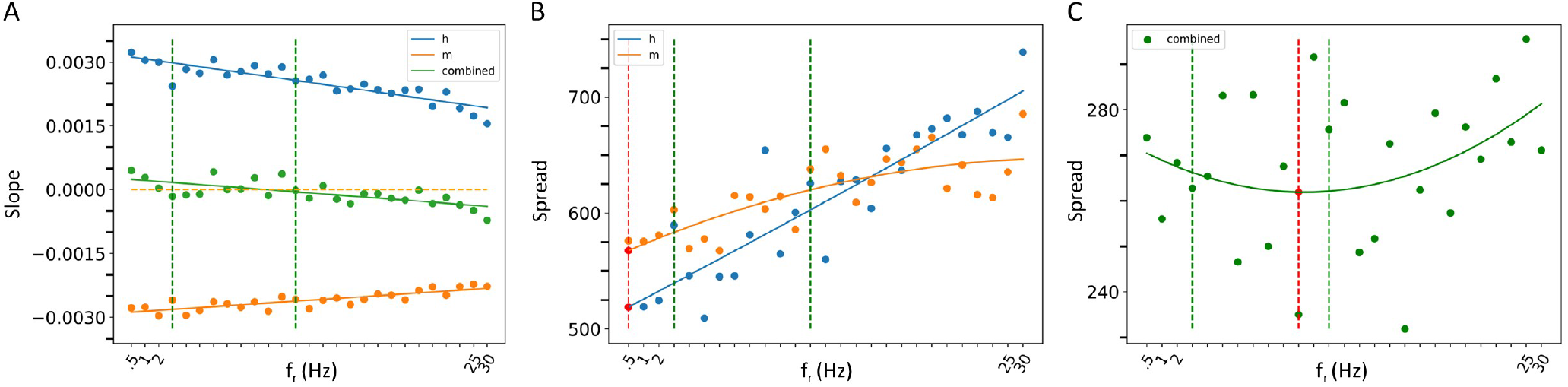
Spread and Slope analysis. **(A)** Slope analysis. The lines are linear fit lines for Ih alone (blue dots), IM alone (orange dots) or IM and Ih combined (green dots). The orange dashed line is where the slope is zero. **(B,C)** Spread analysis. The curves are parbolic fits for Ih alone (blue dots), IM alone (orange dots) or IM and Ih combined (green dots) in (C). The red dashed lines show where the local minima of the parabolic fits are with the minimum shown with a red dot. The theta frequency range (3-12 Hz) is delineated with vertical green dashed lines in all of the plots.

In our spread analysis, we found that as *f_r_* increased, the spread in both IM and Ih STA plots increased as shown in FIGURE 8B. However, when we examined a combination of Ih and IM, the spread was more variable as shown in FIGURE 8C. Parabolic fits to each of IM, Ih and their combination were performed and the curve fits are shown in FIGURE 8B&C. From these fits, it is clear that the combined IM and Ih exhibits a tendency to have a minimal value in the theta frequency range, unlike IM or Ih alone, although none of these parabolic fits were found to be statistically significant. However, one can obtain a statistically significant linear fit to Ih (*R*^2^=83.1%). While the parabolic fits were not statistically significant, a nonlinear local polynomial regression can yield a statistically significant fit with a minimal value for the combined IM and Ih, but neither IM nor Ih alone. Overall, the findings of our spread and slope analysis imply that to specifically have spiking resonance at theta frequencies, we need to have a balance. That is, our analyses indicate that there is a ”balanced sweet spot” during which the combination of Ih and IM minimally change approaching spiking (zero slope and minimal spread), regardless of the baseline firing frequency of the OLM cell *in vivo*. The combination of their biophysical characteristics is what allows this, and not either current on its own.

### 3.3 Parameter estimation with RAUKF

Having now determined that particular biophysical currents contribute to theta frequency spiking resonance, we move to consideration of parameter estimation directly from experimental data. That is, we examine whether we could directly estimate parameter values from experimental recordings using Kalman Filtering (KF) techniques. If so, then it may be possible to bypass the development of the detailed, multi-compartment cell models to get insights into relative differences of biophysical characteristics in various cell types under various conditions. For example, conductance or time constant values of various biophysical channels in a given cell type would vary during different neuromodulatory states to affect spiking output. To be able to do this, we first need a mathematical model structure and a set of parameters whose values are to be estimated. Using a single compartment model is ideal, but not required, in applying KF parameter estimation techniques.

For the OLM cell, we have models of SINGLE and FULL and use them as a proof of concept in doing direct parameter estimations. We carry out two sets of parameter estimations in which we assume that the experimental output is represented by the membrane potential of the model cell generated from SINGLE or FULL. That is, we use either SINGLE or FULL to generate observation data for parameter estimation. In both cases, we focus on estimating parameter values for the same four conductances that were optimized in creating the reduced single compartment model (see Methods). We use SINGLE’s mathematical model structure (see SUPPLEMENTARY Material for equations) and estimate parameter values for *g_M_, g_Kdrf_, g_Ka_, g_Na_* via RAUKF (see Methods for details). The protocol we use is shown in FIGURE 9, and we use initial conditions that are either optimal (i.e., the optimized conductance values from SINGLE) or poor (just the order of magnitude).

**Figure 9.**
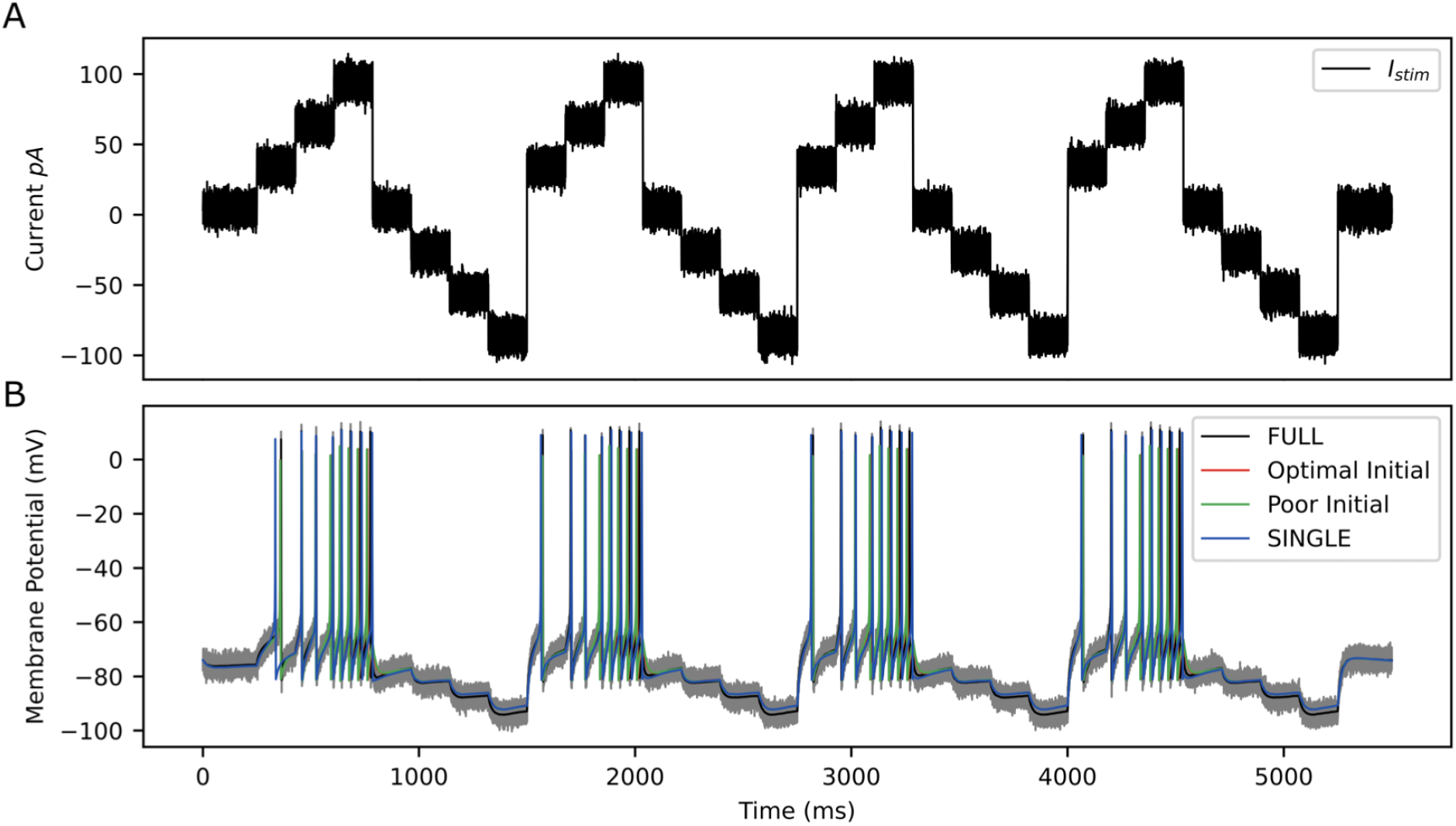
RAUKF input protocol. **(A)** Noisy input protocol applied to mathematical model to obtain parameter estimates. It is a noisy step current (*σ* = 5 *pA*), with a mean step of 30, 60, 90, 0, 30, 60, 90 *pA*, for 196 *ms* each repeated four times beginning with 250 *ms* of noisy current. **(B)** Gray trace is observation used in filter, black trace is the real membrane potential, FULL. Red and green traces are RAUKF model estimated behaviour initialized with parameters determined with optimal and poor initial estimates respectively processed observing the FULL model. Blue trace is model SINGLE, parameters determined by BluePyOpt.

Parameter estimates obtained using the RAUFK algorithm are given in TABLES 2 & 3. If the RAUKF algorithm is working well, then the conductance estimates obtained using SINGLE should be very close to their actual values since in this case the mathematical model structure is the same as what is being used to create the observation data. As shown in TABLE 2, this is clearly the case regardless of the initial conditions used. The parameter estimates are mostly less than one percent different from the actual values. How the algorithm approaches these values is shown in FIGURE 10A.

**Table 2.** Using SINGLE for Experimental Observations

|  | Optimal Initial | Final | % Diff | Poor Initial | Final | % Diff | Actual Value |
| --- | --- | --- | --- | --- | --- | --- | --- |
| $g_M$ | 1.91e-05 | 1.89e-05 | 1.17% | 1.00e-05 | 1.88e-05 | 1.31% | 1.91e-05 |
| $g_{Kdrf}$ | 4.33e-03 | 4.31e-03 | 0.50% | 1.00e-03 | 4.31e-03 | 0.63% | 4.33e-03 |
| $g_{K_a}$ | 7.31e-03 | 7.35e-03 | 0.59% | 1.00e-03 | 7.35e-03 | 0.60% | 7.31e-03 |
| $g_{Na}$ | 4.85e-03 | 4.84e-03 | 0.15% | 1.00e-03 | 4.84e-03 | 0.15% | 4.85e-03 |

**Table 3.** Using FULL for Experimental Observations

|  | Optimal Initial | Final | % Diff | Poor Initial | Final | % Diff | SINGLE Value |
| --- | --- | --- | --- | --- | --- | --- | --- |
| $g_M$ | 1.91e-05 | 1.84e-05 | 3.70% | 1.00e-05 | 1.66e-05 | 12.79% | 1.91e-05 |
| $g_{Kdrf}$ | 4.33e-03 | 5.25e-03 | 21.07% | 1.00e-03 | 5.53e-03 | 27.54% | 4.33e-03 |
| $g_{K_a}$ | 7.31e-03 | 7.17e-03 | 1.87% | 1.00e-03 | 7.15e-03 | 2.19% | 7.31e-03 |
| $g_{Na}$ | 4.85e-03 | 4.55e-03 | 6.06% | 1.00e-03 | 4.56e-03 | 5.90% | 4.85e-03 |

**Figure 10.**
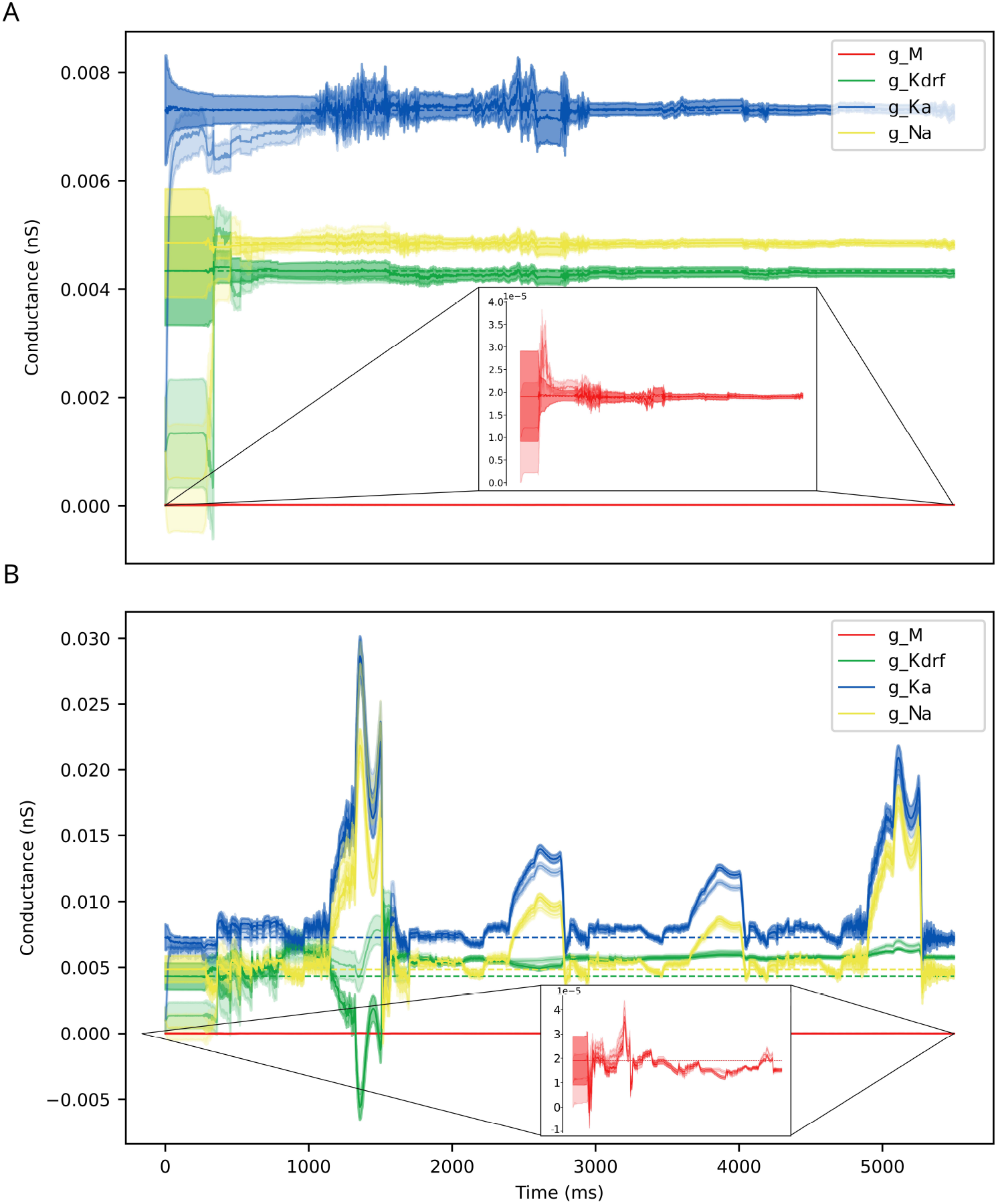
Parameter estimates over time via RAUKF algorithm. Estimate of conductance values over time, shaded region depicts the standard error of the estimate derived from the process covariance **P**. For each conductance, two traces are made, one transparent the other not. The transparent trace of each conductance represents the RAUKF estimates over time when initialized with a poor estimate, the opaque trace depicts conductance estimates with initial values set to the same values as in SINGLE (optimized values). The conductance estimate of *g_M_* is presented in a zoomed frame as the order of magnitude of *g_M_* is 10^−3^ less than the other values. **(A)** RAUKF estimates applied to an observation using SINGLE. **(B)** RAUKF estimates applied to observation using FULL.

TABLE 3 shows the estimated parameters when FULL is used for the observation data (akin to actual experimental data), and how the algorithm approaches these estimated values is shown in FIGURE 10B. The percent differences shown in TABLE 3 are relative to the SINGLE parameter values. This essentially ends up being a comparison with parameter values obtained via the bluepyopt optimization in the reduced model development (see Methods). In the bluepyopt case, the optimized parameter values are obtained using metrics that include spike height, frequency etc. considering three different step current values, whereas in the RAUKF case, a noisy input protocol (FIGURE 9 is used to obtain parameter estimates. As shown in FIGURE 11 for a 60 pA input, while they are not identical, they are very similar. Comparisons for 30 and 90 pA step inputs are shown in SUPPL FIGURES S10 & S11.

**Figure 11.**
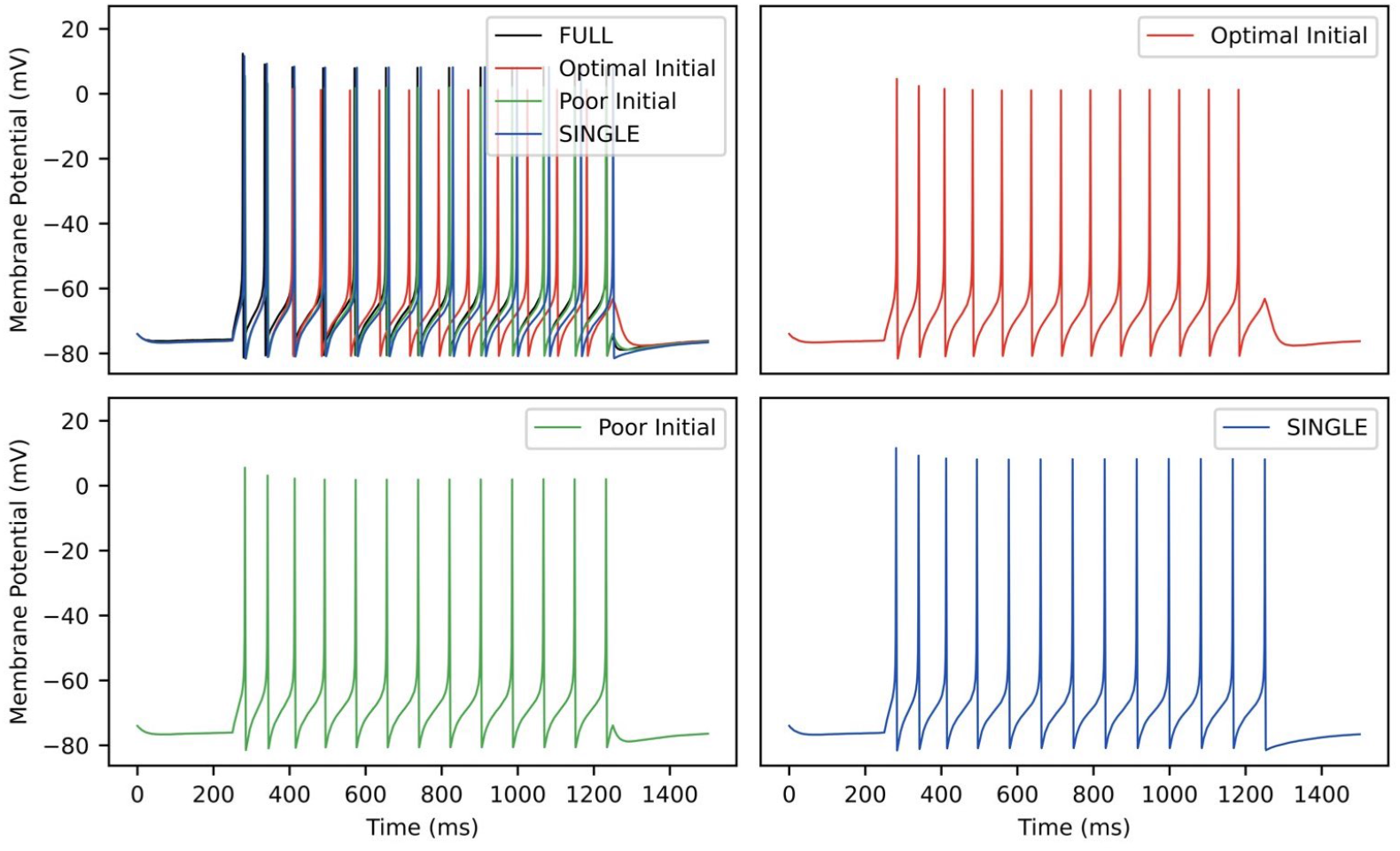
RAUKF Parameter Comparison. Traces of FULL (black), SINGLE (blue) and RAUKF model poor and optimal estimates (green and red) to a step 60 *pA* step current for one second, similar to as shown in FIGURE 1. The traces of SINGLE and RAUKF are overlayed atop the FULL’s in the top left. The remaining corners are the traces of SINGLE and RAUKF separated for comparison.

## 4 DISCUSSION

In this work, we produced a reduced, single compartment model of an OLM cell that has similar biophysical current balances to a previously developed full, multi-compartment OLM cell model that was produced in a ‘neuron-to-model’ matched fashion (Sekulić et al., 2020). Using this reduced model, we showed that it produced similar behaviours to the multi-compartment one regarding phase-dependent contributions of biophysical currents and spiking resonances in *in vitro* and *in vivo*-like states (Guet-McCreight and Skinner, 2021). Due to its reduced nature, we were able to produce a much wider range of firing frequencies for *in vivo*-like states to encompass OLM cell firing ranges observed experimentally. In turn, this allowed us to produce a large dataset of a wider and larger range of spike frequency resonances that could be subsequently explored from underlying biophysical perspectives. Using spike-triggered analyses, we were able to show that characteristics from a combination of M- and h-currents are what allow spiking resonances at theta frequencies specifically to be present. Given the slower time constants of h- and M-currents relative to the other current types, it makes sense that they could play larger roles for theta frequencies in particular. It is further interesting that these channel types could be playing an oversized role in generating theta frequency spiking resonances with their distinct cholinergic profiles and prominent h-channels with location-dependent characteristics (Hilscher et al., 2019; Lawrence, 2008; Maccaferri, 2005). That is, there may be particular ways by which modulatory influences on M- and h-currents of OLM cells could control theta rhythmic activities. It is particularly interesting that we found that it is a *combination* of these two current types that is important, and not just one of them alone, suggesting that modulatory balances need to be in play for the existence of theta frequency spiking resonances.

Using our full, multi-compartment OLM cell model as an experimental proxy, we showed that it is possible to directly estimate parameter values from voltage recordings using a noisy input protocol that used multiple current steps. We limited our examination to estimating parameters of maximal conductances of four channel types but this is not a restriction of the RAUKF algorithm itself (Azzalini et al., 2022). Rather, we sought a proof of principle for the approach since we had both full, detailed multi-compartment OLM cell models and reduced single compartment models, the latter’s mathematical model structure that was used with the RAUKF algorithm. Our choice to focus on extracting conductances *g_M_, g_Kdrf_, g_Ka_* and *g_Na_* also allowed us to directly compare these extracted parameter values with those obtained in developing the reduced model using a genetic optimization technique (BluePyOpt) to match feature values. Although the resulting parameter values were not identical, they were very similar with the most difference being in *g_Kdrf_*. The RAUKF estimate was larger which would explain why the spike height is a bit less (see FIGURE 11). The spike height match using BluePyOpt is expected since spike height was a specific feature that was included during the optimization.

The successful demonstration of the RAUKF algorithm here is strong encouragement to expand its usage to estimate additional parameter values in the OLM cell model and to consider application to other cell types. Indeed, these KF techniques have already been applied in various ways that include single neuron dynamics (Lankarany et al., 2013, 2014; Schiff, 2009; Ullah and Schiff, 2009). A major advantage of using the RAUKF algorithm over other optimization techniques is its speed and possibility for real-time usage. This could be particularly exciting as the modulatory nature of these h- and M-currents could be monitored in real time and changes under different conditions could be assessed.

Obtaining *in vivo* recordings of identified cells are highly non-trivial and it is important to point out how our work has built on and gone beyond previous studies. In earlier studies, artificial *in vivo* firing rates of OLM cells were restricted to 2.5 Hz (Kispersky et al., 2012; Sekulić and Skinner, 2017), but they can be much higher (Katona et al., 2014; Varga et al., 2012). Our recent modelling study that included OLM cells during IVL states produced a restricted range of firing rates using previously optimized synaptic characteristics (Guet-McCreight and Skinner, 2020). Here, with our developed reduced model, we were less restricted as we focused on having valid *in vivo* characteristics and not on optimized synaptic characteristics based on defined presynaptic cell types. The fact that our reduced single compartment model was able to capture complex behaviours to those seen with the full, multi-compartment model suggests that including dendritic OLM cell details are not overly critical for the consideration of spiking resonances, possibly due to the compact nature of OLM cells. Of course, this does not mean that dendritic details and channel distributions are not relevant. Rather, it suggests that a consideration of somatic biophysical balances of h- and M-currents is sufficient to gain insight into theta frequency spiking resonances. Further, all the different channel types present in the multi-compartment model were not retained in the reduced single compartment one. Specifically, calcium channels types were not included as we knew that they could not be biophysically balanced in a single compartment model. One can consider developing a reduced two-compartment model to include these additional biophysical currents in future studies.

In closing, we would like to heartily agree with statements that “diversity is beneficial” to have an “immense impact on our understanding of the brain”, as stated in an excellent, recent review on neural modelling that pushes for combinations and not fragmentation (Eriksson et al., 2022). Our work has used developed biophysically detailed mathematical models based on *in vitro* data, created artificial *in vivo* states with reduced biophysical models to capture the range of firing frequencies *in vivo*, and directly extracted parameter values from voltage recordings of an experimental proxy (i.e., detailed, multi-compartment models). In doing this, we now have an approach and a focus to directly examine and gain insight into theta rhythms in the hippocampus from the perspective of h- and M-currents in OLM cells’ control of theta frequency spiking resonances. Diversity is beneficial!

## AUTHOR CONTRIBUTIONS

All authors contributed to conception and design of the study. ZS performed model simulations and all related analyses, DC performed KF analyses and usage. ZS, DC and FS wrote the first draft of the manuscript. All authors contributed to manuscript revision, read, and approved the submitted version.

## ACKNOWLEDGMENTS

We thank Miguel Barreto for statistical analyses and Alexandre Guet-McCreight for sharing original data points for comparisons and earlier technical assistance.

## SUPPLEMENTAL DATA

Supplementary Material consists of equation details and additional figures and tables.

