## Supplementary Material for "Reduced oriens-lacunosum/moleculare (OLM) cell model identifies biophysical current balances for *in vivo* theta frequency spiking resonance"

#### 1 MODEL BIOPHYSICAL CURRENTS

In this section, we provide equation details and parameter values for the different ion channel types producing the biophysical currents that are present in model FULL in the main text, that is, our previously published detailed multi-compartment OLM cell model (Sekulić et al., 2020). The conductance-based mathematical formulation specifics were adapted from previous OLM cell models (Lawrence et al., 2006; Sekulić et al., 2014).

The different channel types are transient sodium, fast and slow delayed rectifier potassium, A-type potassium, M-type, hyperpolarization-activated cation (h-), T- and L-type calcium, and calcium-dependent potassium channels. The maximal conductance parameters are represented respectively as $G_{NaT}, G_{Kdrf}, G_{Kdrs}, G_{KA}, G_h, G_M, G_{CaT}, G_{CaL}, G_{KCa}$. For channels that have different distribution regarding location, their maximal conductance parameters are denoted with $(s)$, $(a)$ or $(d)$ to indicate soma, axon, or dendrites. Maximal conductance values are given in TABLE S1. Reversal potentials for sodium ($E_{Na}$), potassium ($E_K$) and h-channels ($E_h$) are 90, -95 and -34 mV respectively, and the leak reversal potential ($E_L$) is -64.6 mV. Axial resistivity ($R_a$) is 125.24 $\Omega$ cm and specific capacitance ($C_m$) is 0.27 $\mu F/cm^2$. Temperature scalar of $qT = 3^{(T-23)/10}$ with $T$ =34 degC.

As they are helpful for comments made in the main text, we also include time constant and steady-state activation plots for hyperpolarization-activated cation and M-type channels.

**Table S1.** Location and maximal conductance values for ion channel types.

| **Conductance type** | **Distribution location** | **Conductance value** ($pS/\mu m^2$) |
|---|---|---|
| $G_{NaT,s}$ | soma | 70.986 |
| $G_{NaT,d}$ | dendrites | 99.478 |
| $G_{NaT,a}$ | axon | 66.418 |
| $G_{Kdrf,s}$ | soma | 115.47 |
| $G_{Kdrf,d}$ | dendrites | 50.490 |
| $G_{Kdrf,a}$ | axon | 155.97 |
| $G_{Kdrs,s}$ | soma | 0.0054154 |
| $G_{Kdrs,d}$ | dendrites | 0.0038488 |
| $G_{Kdrs,a}$ | axon | 0.0081732 |
| $G_{KA}$ | soma, dendrites | 76.077 |
| $G_{CaL}$ | dendrites | 47.187 |
| $G_{CaT}$ | dendrites | 1.0113 |
| $G_{KCa}$ | dendrites | 1.8194 |
| $G_M$ | soma, dendrites | 0.13738 |
| $G_h$ | soma, dendrites | 0.10631 |
| $G_L$ | soma, dendrites | 0.075833 |

**Passive leak current:** $I_L = G_L\ (V - E_L)$

**Transient sodium current:** $I_{Na} = G_{NaT} \cdot m^3 h(V - E_{Na})$

$$\frac{dm}{dt} = \alpha_m(1-m) - \beta_m m \quad \text{(S1)}$$

$$\frac{dh}{dt} = \alpha_h(1-h) - \beta_h h \quad \text{(S2)}$$

where, for somatic compartments,

$$\alpha_m(V) = \frac{-0.1(V + 38 - V_{shift})}{\exp(-(V + 38 - V_{shift})/10) - 1} \quad \text{(S3)}$$

$$\beta_m(V) = 4\exp(-(V + 63 - V_{shift})/18) \quad \text{(S4)}$$

$$\alpha_h(V) = 0.07\exp(-(V + 63 - V_{shift})/20) \quad \text{(S5)}$$

$$\beta_h(V) = \frac{1}{1 + \exp(-(V + 33 - V_{shift})/10)} \quad \text{(S6)}$$

$$V_{shift} = -4.830346371483079\ mV \quad \text{(S7)}$$

and for dendritic and axonal compartments,

$$\alpha_m(V) = \frac{-0.1(V + 45 - V_{shift})}{\exp(-(V + 45 - V_{shift})/10) - 1} \quad \text{(S8)}$$

$$\beta_m(V) = 4\exp(-(V + 70 - V_{shift})/18) \quad \text{(S9)}$$

$$\alpha_h(V) = 0.07\exp(-(V + 70 - V_{shift})/20) \quad \text{(S10)}$$

$$\beta_h(V) = \frac{1}{1 + \exp(-(V + 40 - V_{shift})/10)} \quad \text{(S11)}$$

$$V_{shift,dend} = 4.846190532969488\ mV \quad \text{(S12)}$$

$$V_{shift,axon} = 2.4859651361041872\ mV \quad \text{(S13)}$$

**Hyperpolarization-activated mixed cation current:** $I_h = G_h \cdot r\,(V - E_h)$

$$\frac{dr}{dt} = \frac{r_\infty - r}{\tau_h} \tag{S14}$$

$$r_\infty = \frac{1}{1 + \exp\left(\frac{V - V_{1/2}}{k}\right)} \tag{S15}$$

$$\tau_h(V) = \frac{1}{\exp(-t_1 - t_2 V) + \exp(-t_3 + t_4 V)} + t_5 \tag{S16}$$

Other parameter values given in TABLE S2, and activation time constant and activation curve plotted in FIGURE S1.

**Table S2.** Parameter values of $\tau_h(V)$

| $t_1$ | $t_2$ | $t_3$ | $t_4$ | $t_5$ | $k\ (mV)$ | $V_{1/2}\ (mV)$ |
|---|---|---|---|---|---|---|
| 8.5657797 | 0.0296317 | -6.9145 | 0.1803 | $4.3566601 \times 10^{-5}$ | 9.9995804 | -103.69 |

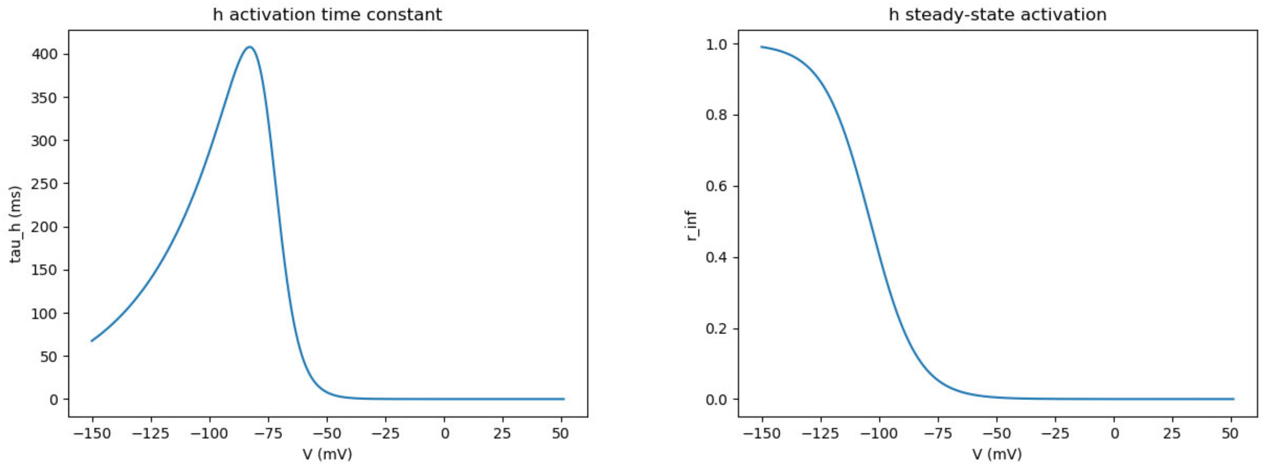


Figure S1: **Time constant and steady state activation of $I_h$.**

**Fast potassium delayed rectifier current:** $I_{Kdrf} = G_{Kdrf} \cdot mh\ (V - E_K)$

$$\tau_m(V)\frac{dm}{dt} = m_\infty - m \quad \text{(S17)}$$

$$\tau_h(V)\frac{dh}{dt} = h_\infty - h \quad \text{(S18)}$$

$$m_\infty(V) = (\frac{1}{1+exp[-(V+36.2)/16.1]})^4 \quad \text{(S19)}$$

$$\tau_m(V) = \frac{27.8exp[(V+33)/14.3]}{qt(1+exp[(V+33)/10])} \quad \text{(S20)}$$

$$h_\infty(V) = \frac{0.92}{1+exp[(V+40.6)/7.8]} + 0.08 \quad \text{(S21)}$$

$$\tau_h(V) = 1000 \quad \text{(S22)}$$

**Slow potassium delayed rectifier current:** $I_{Kdrs} = G_{Kdrs} \cdot mh\ (V - E_K)$

$$\tau_m(V)\frac{dm}{dt} = m_\infty - m \quad \text{(S23)}$$

$$\tau_h(V)\frac{dh}{dt} = h_\infty - h \quad \text{(S24)}$$

$$m_\infty(V) = (\frac{1}{1+exp[-(V+41.9)/23.1]})^4 \quad \text{(S25)}$$

$$\tau_m(V) = \frac{66.7exp[(V+25)/13.3]}{qt(1+exp[(V+25)/6.7)]} \quad \text{(S26)}$$

$$h_\infty(V) = \frac{0.93}{1+exp[(V+52.2)/15.2]} + 0.07 \quad \text{(S27)}$$

$$\tau_h(V) = 1000 \quad \text{(S28)}$$

**A-type potassium transient current:** $I_{Ka} = G_{KA} \cdot mh\ (V - E_K)$

$$\tau_m(V)\frac{dm}{dt} = m_\infty - m \quad \text{(S29)}$$

$$\tau_h(V)\frac{dh}{dt} = h_\infty - h \quad \text{(S30)}$$

$$m_\infty(V) = (\frac{1}{1+exp[-(V+41.4)/26.6]})^4 \quad \text{(S31)}$$

$$\tau_m(V) = 0.5/qT \quad \text{(S32)}$$

$$h_\infty(V) = \frac{1}{1+exp[(V+78.5)/6]} \quad \text{(S33)}$$

$$\tau_h(V) = 0.17(V+105)/qt \quad \text{(S34)}$$

**Muscarinic channel current:** $I_M = G_M \cdot m\ (V - E_K)$

$$\tau_m(V)\frac{dm}{dt} = m_\infty - m \tag{S35}$$

$$m_\infty(V) = \frac{1}{1 + exp[-(V+27)/7]} \tag{S36}$$

$$\tau_m(V) = \frac{1}{0.003/exp[-(V+63)/15] + 0.003/exp[(V+63)/15]} \tag{S37}$$

Activation time constant and activation curve plotted in FIGURE S2.

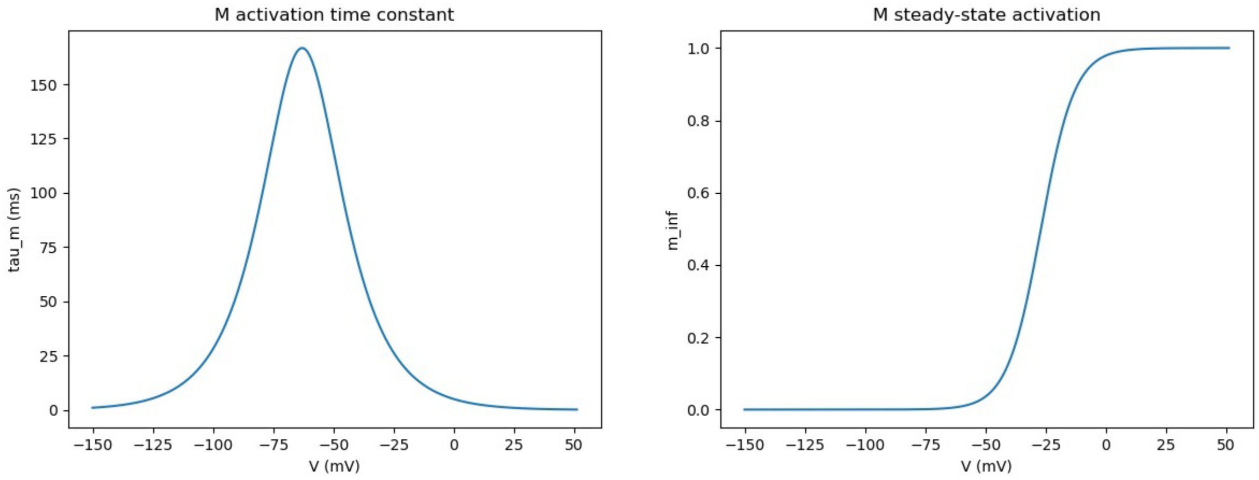


Figure S2: **Time constant and steady state activation of** $I_M$.

**L-type calcium channel current:** $I_{CaL} = G_{CaL} \cdot m^2 h \, \Gamma(V, [Ca^{2+}]_i, [Ca^{2+}]_o)$

$$\Gamma(V, [Ca^{2+}]_i, [Ca^{2+}]_o) = -k(1 - \frac{[Ca^{2+}]_i}{[Ca^{2+}]_o} exp(\frac{V}{k}) \frac{V}{k(exp(\frac{v}{k}) - 1)}) \tag{S38}$$

$$k = 0.043T + 11.65 \tag{S39}$$

$$\frac{dm}{dt} = \alpha_m(1 - m) - \beta_m m \tag{S40}$$

$$\alpha_m(V) = \frac{15.69(-(V - 81.5))}{exp(-(V - 81.5)/10) - 1} \tag{S41}$$

$$\beta_m(V) = 0.29 \exp(-V/10.86) \tag{S42}$$

$$h = \frac{0.001}{0.001 + [Ca^{2+}]_i} \tag{S43}$$

**T-type calcium channel current:** $I_{CaT} = G_{CaT} \cdot m^2 h \, ([Ca^{2+}]) \, \Gamma(V, [Ca^{2+}]_i, [Ca^{2+}]_o)$

$$\Gamma(V, [Ca^{2+}]_i, [Ca^{2+}]_o) = -k(1 - \frac{[Ca^{2+}]_i}{[Ca^{2+}]_o} exp(\frac{V}{k}) \frac{V}{k(exp(\frac{v}{k}) - 1)}) \tag{S44}$$

$$k = 0.043T + 11.65 \tag{S45}$$

$$\frac{dm}{dt} = \alpha_m(1 - m) - \beta_m m \tag{S46}$$

$$\frac{dh}{dt} = \alpha_h(1 - h) - \beta_h h \tag{S47}$$

$$\alpha_m(V) = \frac{0.2(-(V - 19.26))}{exp(-(V - 19.26)/10) - 1} \tag{S48}$$

$$\beta_m(V) = 0.009 \exp(-V/22.03) \tag{S49}$$

$$\alpha_h(V) = 1 \times 10^{-6} exp(-V/16.26) \tag{S50}$$

$$\beta_h(V) = \frac{1}{exp(-(V - 29.79)/10) + 1} \tag{S51}$$

**Calcium-activated potassium current:** $I_{KCa} = G_{KCa} \cdot m([Ca^{2+}]_i) \, (V - E_K)$

$$\frac{dm}{dt} = \alpha_m(1 - m) - \beta_m m \tag{S52}$$

$$\alpha_m(V, [Ca^{2+}]_i) = \frac{0.28[Ca^{2+}]_i}{[Ca^{2+}]_i + 4.8 \times 10^{-4} exp(-0.2V/(273.15 + T))} \tag{S53}$$

$$\beta_m(V, [Ca^{2+}]_i) = \frac{0.48}{1 + [Ca^{2+}]_i/[1.3 \times 10^{-7} exp(-0.24V/(273.15 + T))]} \tag{S54}$$

Calcium concentrations in L-type and T-type calcium and calcium-activated potassium currents are given in the subsection below.

#### 1.1 Calcium dynamics

The original detailed multi-compartment model of Sekulić et al. (2020) is slightly adjusted as detailed below to create model FULL in the main text.

In essence, the calcium modeling is modified in consideration of Migliore et al. (1995).
Model DB link: `https://senselab.med.yale.edu/modeldb/ShowModel?model=3263&file=/ca3_db/cadiv.mod#tabs-1`. The model assumes that the cross-section of each segment is a circle. It models concentration of calcium by dividing this circle into four concentric shells (starting at 0 from the out-most shell). The inner three shells have the same thickness while shell 0 has half of that thickness. $[Ca]_i$ is defined as the concentration of calcium in shell 0. There are four mechanisms that manipulate the calcium ions in the cell. First is calcium influx that translates calcium current into the amount of calcium ions that enter shell 0. Second is calcium diffusion that happens across adjacent shells. Third is calcium buffering within each shell. Fourth is a membrane-bound calcium pump that moves calcium in shell 0 to the extracellular space.

In our modifed version, instead of four shells, we use only one shell, which is the region defined from the membrane to 1/6 diameter deep. Just like Migliore et al. (1995)'s model, there is calcium influx due to calcium currents, buffering within the shell, and a calcium pump that removes free calcium from the cell, but there is not diffusion across shells since there is only one shell.

The following equation describes how calcium current is translated to calcium influx (in moles/time) to shell 0: $Ca_{influx} = -I_{Ca}/(2 \cdot F) * surface\ area$ where $F$ is Faraday's constant.

The following equations describe the kinetics of the membrane-bound calcium pump:

$$\text{Ca}_i + \text{pump} \xleftrightarrow[k_2]{k_1} \text{PumpCa}$$
$$\text{pumpCa} \xleftrightarrow[k_4]{k_3} \text{pump} + \text{Ca}_o$$

$$[pump] + [pumpCa] = 0.2mol/cm^2 \quad \text{(S55)}$$
$$k_1 = 1 \times 10^{10} \mu m^3/ms \quad \text{(S56)}$$
$$k_2 = 5 \times 10^8 ms^{-1} \quad \text{(S57)}$$
$$k_3 = 1 \times 10^{10} ms^{-1} \quad \text{(S58)}$$
$$k_4 = 5 \times 10^6 \mu m^3/ms \quad \text{(S59)}$$
$$\quad \text{(S60)}$$

$$[Ca]'_{i\ pump} = -[Ca]_i[pump] \cdot k_1 + [pumpCa] \cdot k_2 \quad \text{(S61)}$$
$$[pump]' = -[Ca]_i[pump] \cdot k_1 + [pumpCa] \cdot k_2 + [pumpCa] \cdot k_3 \quad \text{(S62)}$$
$$[pumpCa]' = -[pumpCa] \cdot k_2 - [pumpCa] \cdot k_3 + [Ca]_i[pupmp] \cdot k_1 + [pump][Ca]_i \cdot k_4 \quad \text{(S63)}$$
$$[Ca]_o = 2mM \quad \text{(S64)}$$

The following equations describe calcium buffering within the shell:

$$\mathrm{Ca}_i + \mathrm{Buffer} \underset{k_b}{\overset{k_f}{\longleftrightarrow}} \mathrm{CaBuffer}$$

$$k_f = 500mM^{-1}ms^{-1} \quad \text{(S65)}$$

$$k_b = 0.5ms^{-1} \quad \text{(S66)}$$

$$[Ca]'_{buffering} = -[Ca]_i[Buffer] \cdot k_f + [CaBuffer] \cdot k_b \quad \text{(S67)}$$

$$[Buffer]'_{shell\ i} = [Ca]'_{buffering} \quad \text{(S68)}$$

$$[CaBuffer]' = [CaBuffer] \cdot f_b \quad \text{(S69)}$$

From all the above equations, the change of $[Ca]_i$ is as follows:

$$[Ca]'_i = \frac{Ca_{influx}}{Volume_{shell}} + [Ca]'_{buffering} + [Ca]'_{i\ pump} \quad \text{(S70)}$$

#### 2 SINGLE COMPARTMENT OLM CELL MODEL DEVELOPMENT

SINGLE is a single compartment conductance model that maintains some of the currents of FULL.

As noted in the main text, *Cell 1 NMM model* was directly matched with experimental data in which Ih characteristics were obtained in the same cell as passive properties, and thus, we did not try to optimize these aspects directly. The surface area was slightly modified (about 7% difference) from that of FULL (soma and dendrites) to be 31,403.4 $\mu m^2$, and specific capacitance and reversal potentials were kept the same as FULL/original model. The total Ih is 3.34 nS (6.5% difference from original model), with input resistance of 419.9 M$\Omega$ (about 2% difference) and time constant of 34.6 mS (about 5% difference). As a preliminary step, we hand-tuned the maximal conductances of the other currents (INa, IKdrf, IKa, IM) in a single compartment model to test the feasibility of replicating behavior from FULL using a single compartment and fewer current types. These values were then used as initial values in doing an optimization.

We refer to the maximal conductances of IKa, IKdrf, IM and INa with small g's to distinguish them from FULL. They were optimized using BluePyOpt (Van Geit et al., 2016) based on comparison with FULL. The types of features considered were exactly the same as those used in Sekulić et al. (2020). We extracted feature values from the voltage traces at the soma of FULL using 30, 60, and 90 pA current injections. The DEAP optimizer was used to run for 200 generations, with 100 off-springs each generation, 0.15 mutation rate, and 0.85 crossover probability. These parameters were also taken from Sekulić et al. (2020). Initial and optimized values are given in TABLE S3. Voltage trace comparisons of SINGLE and FULL at 30, 60 and 90 pA steps are shown in FIGURE S3. Additional FIGURE S4 shows the currentscapes with the voltage firing for -120, 30 and 90 pA for FULL and SINGLE. For the -120 pA step, we note that the obvious currentscape difference for IKa is due to somatic and dendritic aspects. That is, there are no dendrites for SINGLE. Otherwise, a balance of currents can be seen, similar to 60 pA step that is shown in the main text. The feature differences are shown in FIGURE S5, and the numbered features are described in TABLE S4.

Voltage output of the single compartment model is given by:

$$C_m \frac{dV}{dt} = -I_{Na} - I_{Kdrf} - I_{Ka} - I_M - I_h - I_L + I_{stim} \tag{S71}$$

where biophysical current equations ($I_{Na}, I_{Kdrf}, I_{Ka}, I_M, I_h, I_L$ are given above and the somatic formulation is used for $I_{Na}$. $I_{stim}$ is used when replicating experimental protocols - see Sekulić et al. (2020). Non-optimized values of $g_h$ and $g_L$ are 0.106309 and 0.075833 $pS/\mu m^2$ respectively.

**Table S3.** Initial and optimized conductance values.

| **Conductance type** | **Initial value** ($pS/\mu m^2$) | **Optimized value** ($pS/\mu m^2$) |
|---|---|---|
| $g_{Na}$ | 70.986 | 48.472 |
| $g_{Kdrf}$ | 115.47 | 43.342 |
| $g_{Ka}$ | 76.08 | 73.062 |
| $g_M$ | 0.137 | 0.19082 |

#### 2.1 Additional Supplementary Figures

The IVL spike resonance to directly compare SINGLE and FULL is shown in FIGURE S6.

FIGURE S7 is an additional figure that shows the underlying distribution of parameters for IVL states.

Analogous to FIGURE 7 in the main text, we show STA plots. However, in FIGURE S8 here we show STA plots ranging from -180 to -10 ms (instead of -195 to 25 ms as in main text) so that 'curving up' can be seen.

In FIGURE S9, analogous to FIGURE 7 in the main text, we show normalized currents and means and standard deviation but for a given $f_r$ of 5 Hz and for IM and Ih combined. The reduction in spread for the combined currents relative to the individual ones can be seen.

### 3 ROBUST ADAPTIVE KALMAN FILTER (RAUKF) DETAILS

The equations detailing the states ($\mathbf{x}_k$ and covariance ($\mathbf{Q}$, $\mathbf{R}$) and their respective equations are described within this section in some detail, as they were updated to produce the estimates in the main text.

$$\mathbf{v}_{k|k-1} = \mathbf{y}_k - \mathbf{g}(\hat{\mathbf{X}}_{k|k-1}) \tag{S72}$$

$$\begin{aligned}\hat{\mathbf{Q}}_{k-1} &= \mathbb{E}[\hat{\mathbf{M}}_{k-1}\hat{\mathbf{M}}_{k-1}^T] \\ &= \mathbb{E}[(\mathbf{K}_k\mathbf{v}_{k|k-1})(\mathbf{K}_k\mathbf{v}_{k|k-1})^T] \\ &= \mathbf{K}_k\mathbb{E}[\mathbf{v}_{k|k-1}\mathbf{v}_{k|k-1}^T]\mathbf{K}_k^T \\ &\approx \mathbf{K}_k\mathbf{v}_{k|k-1}\mathbf{v}_{k|k-1}^T\mathbf{K}_k^T\end{aligned} \tag{S73}$$

$$\mathbf{Q}_{k-1} \leftarrow (1-\lambda)\mathbf{Q}_{k-1} + \lambda\hat{\mathbf{Q}}_{k-1} \tag{S74}$$

$$\mathbf{v}_{k|k} = \mathbf{y}_k - \mathbf{g}(\hat{\mathbf{X}}_{k|k}) \tag{S75}$$

$$\begin{aligned}\hat{\mathbf{R}}_k &= \mathbb{E}[\mathbf{v}_{k|k}\mathbf{v}_{k|k}^T] + \hat{\mathbf{P}}_{k|k}^{yy} \\ &\approx \mathbf{v}_{k|k}\mathbf{v}_{k|k}^T + \hat{\mathbf{P}}_{k|k}^{yy}\end{aligned} \tag{S76}$$

$$\mathbf{R}_k = (1-\delta)\mathbf{R}_k + \delta\hat{\mathbf{R}}_k \tag{S77}$$

A simple fault detection rule follows from innovation-based methods and the statistical function

$$\phi_k = \mathbf{v}_{k|k}^T(\hat{\mathbf{P}}_{k|k}^{yy})^{-1}\mathbf{v}_{k|k} \tag{S78}$$

which has $\chi^2$ distribution with $s = 1$ degrees of freedom since our observation is one dimensional (Hajiyev and Caliskan, 2003; Zheng et al., 2018). The innovation vector ($\mathbf{v}_{k|k}$) is normally distributed under normal conditions $\mathcal{H}_0$. If the distribution deviates from a normal distribution a mismatch has occurred $\mathcal{H}_1$, and as such we update $\mathbf{Q}$ and $\mathbf{R}$ to try to correct for the mismatch. To test this difference we used a chi-squared test to determine when $P(\chi^2 > \chi^2_{\alpha,s}) = \alpha$, where $\alpha$ is the significance level and $\chi^2_{\alpha,s}$ denotes the threshold to be exceeded for a fault to be detected (Hajiyev and Soken, 2014):

$$
\begin{aligned}
\mathcal{H}_0: &\quad \phi_k \leq \chi^2_{\alpha,s} \\
\mathcal{H}_1: &\quad \phi_k > \chi^2_{\alpha,s}
\end{aligned}
\tag{S79}
$$

When $\mathcal{H}_0$ is rejected the weights $\lambda$ and $\delta$ are set according to the innovation statistics as well as application dependent parameters $a$ and $b$ in order to update $\mathbf{Q}$ and $\mathbf{R}$ according to (S74) and (S77).

$$
\lambda = \max\left(\lambda_0, \frac{\phi_k - a\chi^2_{\alpha,s}}{\phi_k}\right) \tag{S80}
$$

$$
\delta = \max\left(\delta_0, \frac{\phi_k - b\chi^2_{\alpha,s}}{\phi_k}\right) \tag{S81}
$$

SINGLE, as originally implemented in NEURON, has some slight modifications to the equations described in S1-S54.

The modified equations as used in the NEURON implementation, as well as the Python implementation are depicted below.

$$
m_{\tau|M}(V) = \min\left(\frac{1}{c_1 \mathrm{e}^{\frac{V-V_{\frac{1}{2},1}}{k_1}} + c_2 \mathrm{e}^{-\frac{V-V_{\frac{1}{2},2}}{k_2}}}, 7\right) \tag{S82}
$$

$$
h_{\tau|Ka}(V) = \min\left(\alpha_h \cdot \frac{V - V_{\frac{1}{2},h}}{qt}, 5\right) \tag{S83}
$$

$$
V_{trap}(x, y) = \begin{cases} y \cdot (1 - \frac{x}{2y}) & \frac{x}{y} < 1e-6 \\ \frac{x}{\mathrm{e}^{\frac{x}{y}}-1} & \frac{x}{y} \geq 1e-6 \end{cases} \tag{S84}
$$

$$
m_{\tau|Na}(V) = \frac{1}{0.1 \cdot V_{trap}(-(V + 38 - V_{Shift}), 10)} + 4\mathrm{e}^{-\frac{V+63-V_{Shift}}{18}} \tag{S85}
$$

$$
m_{\infty|Na}(V) = \frac{0.1 \cdot V_{trap}(-(V + 38 - V_{Shift}), 10)}{0.1 \cdot V_{trap}(-(V + 38 - V_{Shift}), 10) + 4\mathrm{e}^{-\frac{V+63-V_{Shift}}{18}}} \tag{S86}
$$

#### 3.1 RAUKF Extended Figures

The stimulation protocol used for the RAUKF procedure was generated to encapsulate responses to low, high, positive and negative currents (see FIGURE 1 in the main text). The performance of the RAUKF estimates were qualitatively compared to SINGLE using different input currents. Output for 30 and 90 pA steps are shown in FIGURES S10 & S11 below, and the 60 pA current step is shown in the main text.

As noted in the Results of the main text, difference in peak height is attributable to the change in $g_{Kdrf}$ estimation. Qualitiatively the spike timing of the RAUKF estimates seem to match better overall to FULL when compared to the BluePyOpt SINGLE estimates. This is particularly noticeable at the end of the step current in the bottom most figure where the RAUKF (poor initial) estimate matches the timing of the polarization of FULL, which is insufficient for a spike.

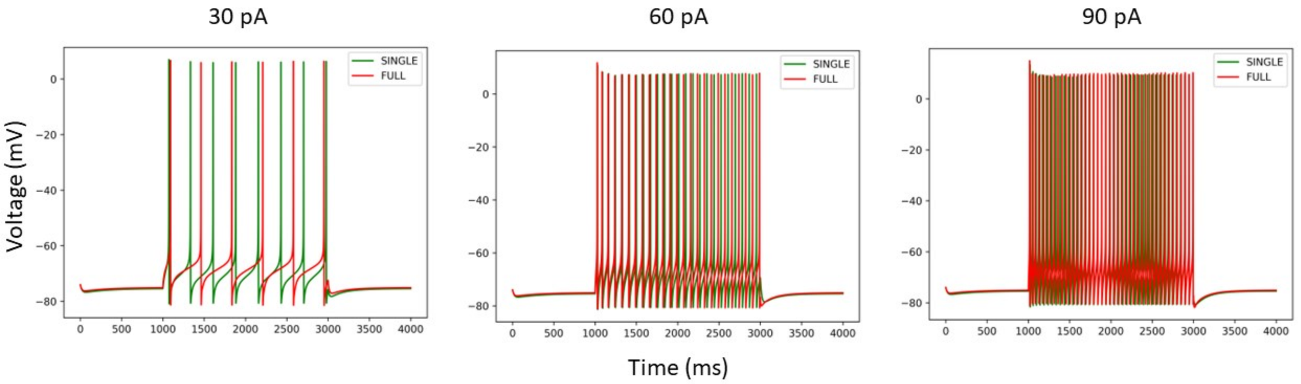


Figure S3: ***In vitro* voltage comparison of SINGLE and FULL.**

**Table S4. Descriptions of the eFEL measurements and the chosen standard deviation values ($\sigma$) that were used as objective features and weights in the optimizations.**

| ***Name*** | $\sigma$ | ***Description*** |
|---|---|---|
| **1.** AP_amplitude_from_voltagebase (mV) | 0.1 | The height of the AP measured from voltage base. |
| **2.** AP_width (ms) | 0.01 | Width of each peak at the value of threshold. |
| **3.** AP_amplitude (mV) | 0.1 | The relative height of the AP between the peak voltage and the voltage where the first derivative is higher than 12 V/s for at least 5 points. |
| **4.** AHP_time_from_peak (ms) | 0.1 | Time between AP peaks and AHP depths. |
| **5.** time_to_first_spike (ms) | 1 | Time from the start of the stimulus to the maximum of the first peak. |
| **6.** voltage_base (mV) | 0.1 | The resting membrane potential before the current step. |
| **7.** AP_amplitude_change | 0.001 | Difference of the amplitudes of the second and the first AP divided by the amplitude of the first AP. |
| **8.** AP_duration_half_width (ms) | 0.01 | Full width at half maximum of each action potential. |
| **9.** AHP_depth (mV) | 0.1 | Relative voltage difference between the minimum AHP voltage and the voltage base. |
| **10.** mean_frequency (Hz) | 0.1 | The mean frequency of the firing rate. |
| **11.** AHP_slow_time | 0.001 | Time difference between absolute voltage values at the first after-hyperpolarization starting 5 ms after the peak and the peak, divided by interspike interval. |
| **12.** adaptation_index | 0.001 | Normalized average difference of two consecutive ISIs. |

AP: Action Potential; AHP: After-Spike Hyperpolarization. Consult the eFEL manual for more details on these measurements: `https://media.readthedocs.org/pdf/efel/latest/efel.pdf`

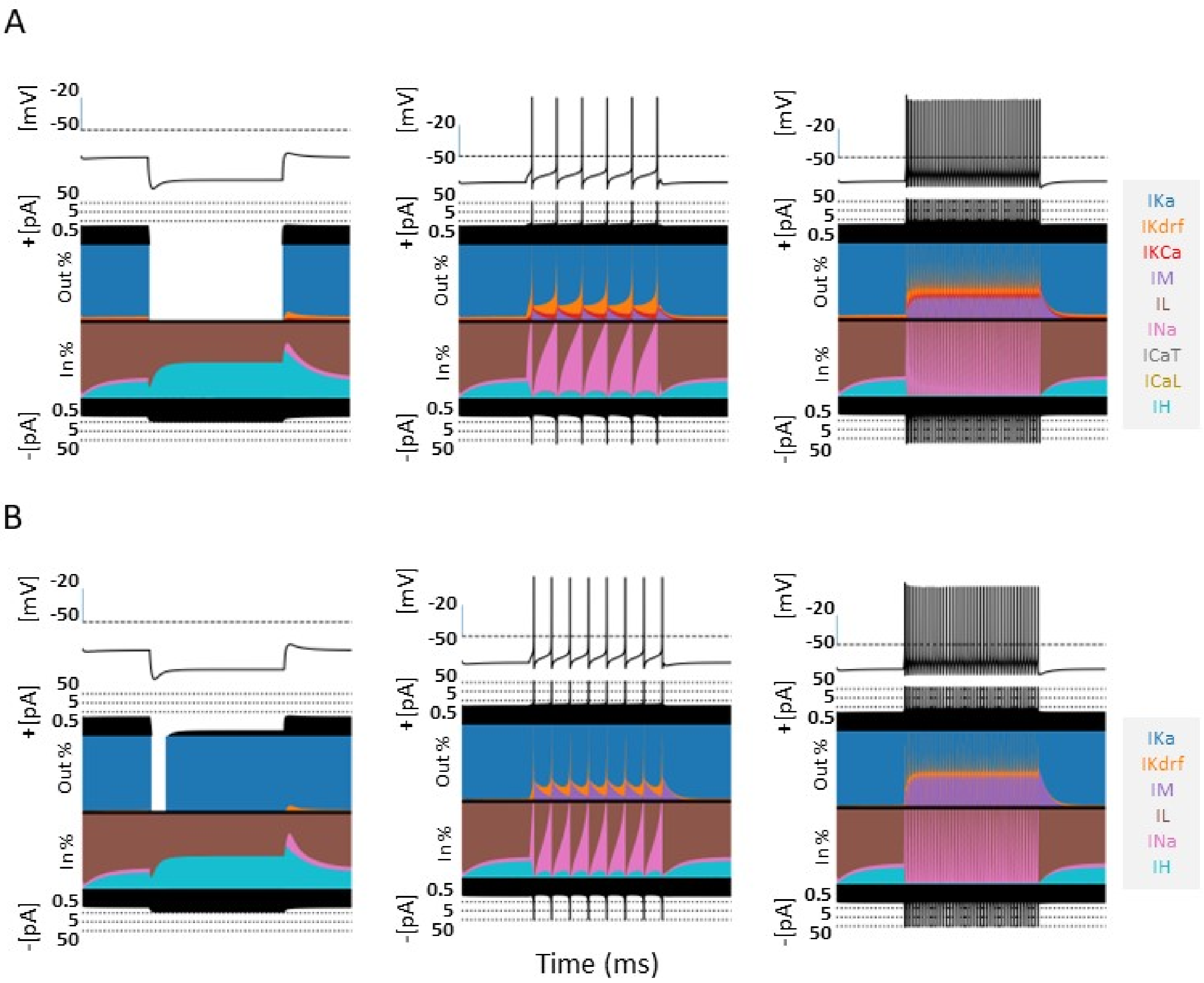


Figure S4: **Comparison of models - FULL and SINGLE.**
Currentscapes are shown for models of FULL (somatic) **(A)** and SINGLE **(B)**. Their firings and corresponding biophysical currents are similar. Simulations were conducted for 4000 ms, with a holding current of 4 pA throughout. From left to right, step currents of -120 pA, 30 pA, and 90 pA were injected respectively at 1000 ms for 2000 ms. 60 pA step is shown in the main text.

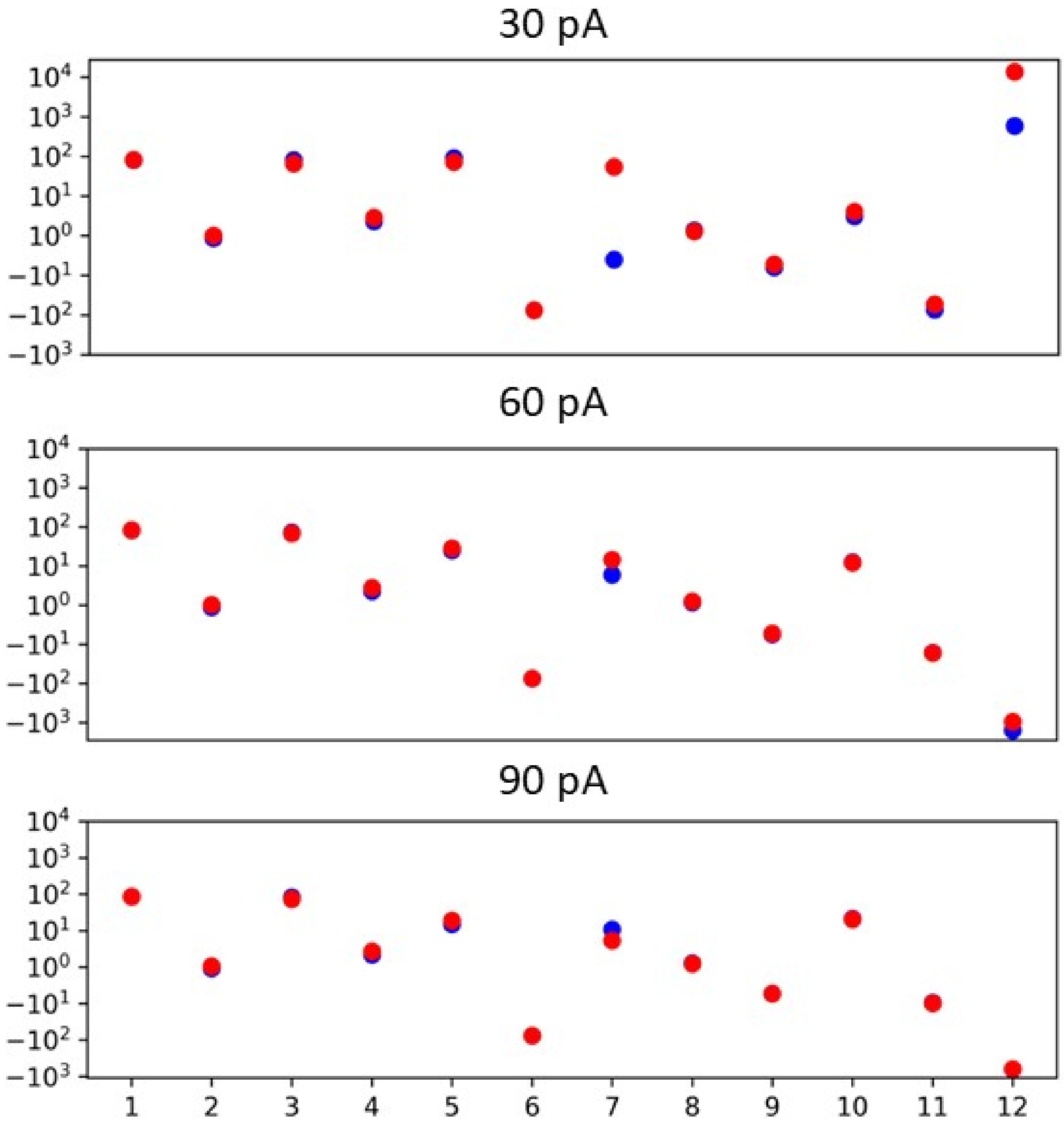


Figure S5: **Measurements of objective e-features.** Optimized values are shown in red and target values are shown in blue. e-feature names and descriptions are given in TABLE S4.

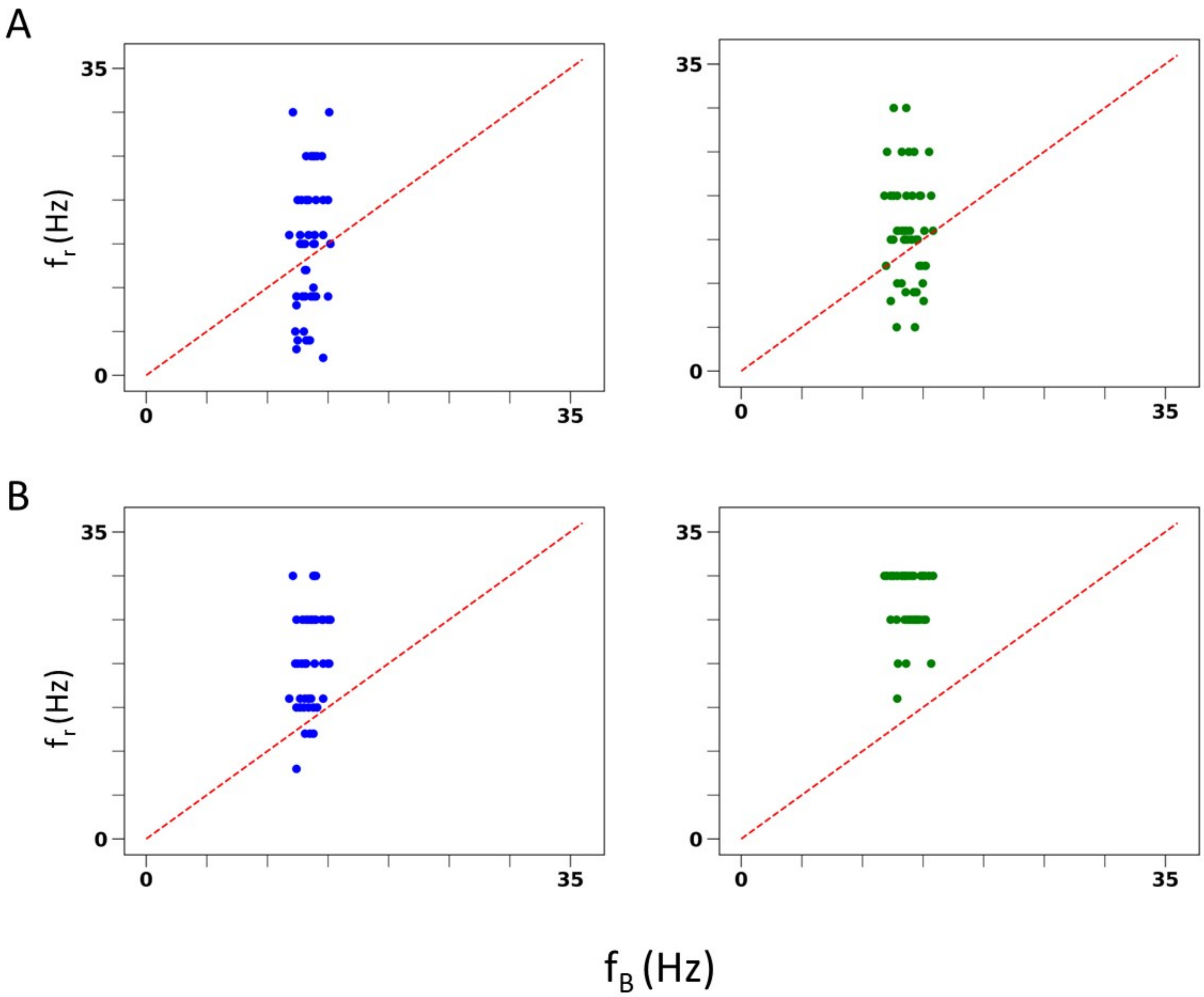


Figure S6: ***IVL* spiking resonance pattern in SINGLE is comparable to FULL. (A)** Resonant frequency ($f_r$) of inhibitory perturbation at different baseline firing rates ($f_B$). Blue dots are from SINGLE, green dots from FULL. **(B)** Resonant frequency ($f_r$) of excitatory perturbation at different baseline firing rates ($f_B$). Blue dots are from SINGLE, green dots from FULL.

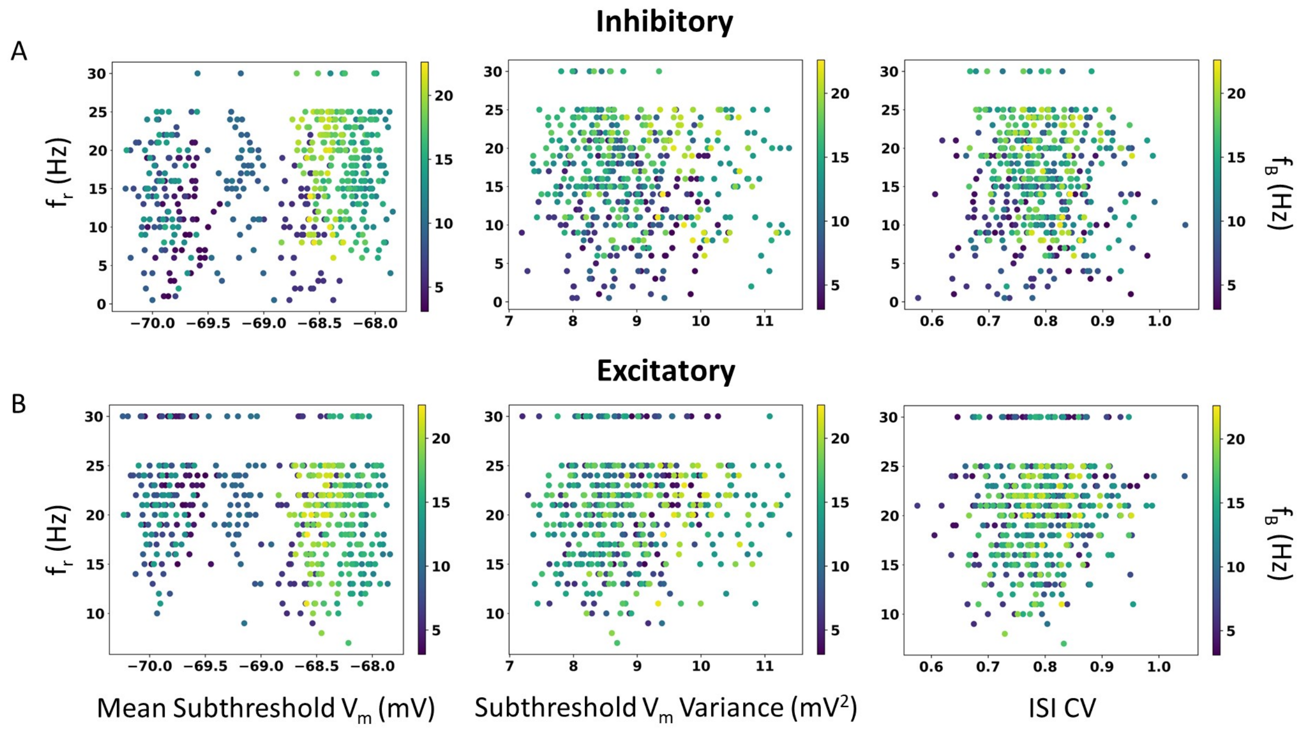


Figure S7: **IVL state expression.** Mean ($\overline{V_m}$) and variance ($\sigma^2_{V_m}$) of the sub-threshold membrane potential, and the $ISICV$ are plotted. $f_B$ are represented by dots colored according to the color bar. As for FIGURE 6 in main text, 500 IVL states are used for illustration to allow individual dots to be visible. Note that for any given $f_r$, there are a range of $\overline{V_m}$, $\sigma^2_{V_m}$ and $ISICV$ values. **(A)** Using inhibitory perturbations. **(B)** Using excitatory perturbations.

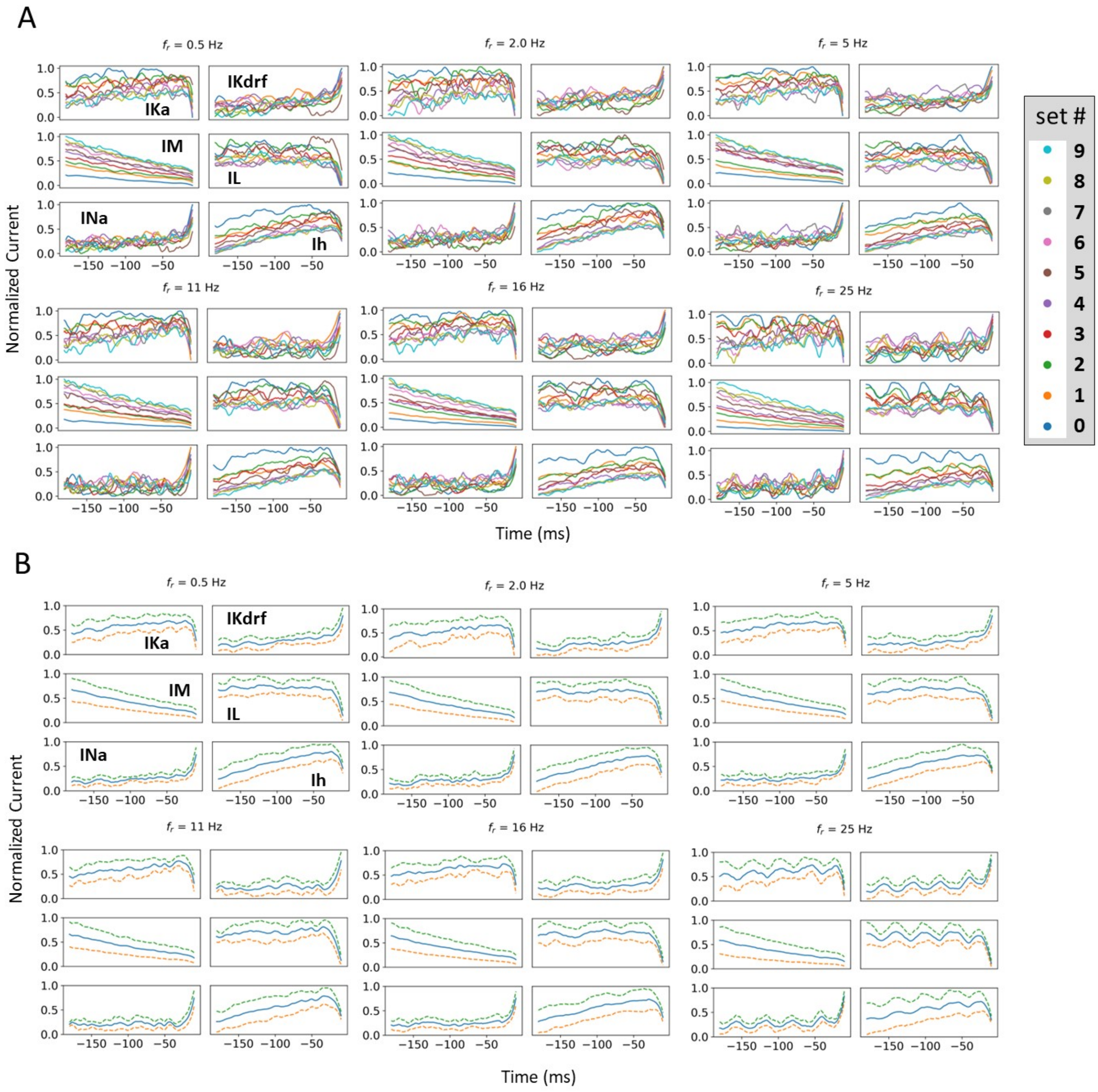


Figure S8: **STAs - different time range.** Exact same layout and examples as FIGURE 7 in the main text with normalized currents **(A)** and means and standard deviations **(B)**, but the range shown is from -180 to -10 ms. In the main text, it is from -195 to -25 ms.

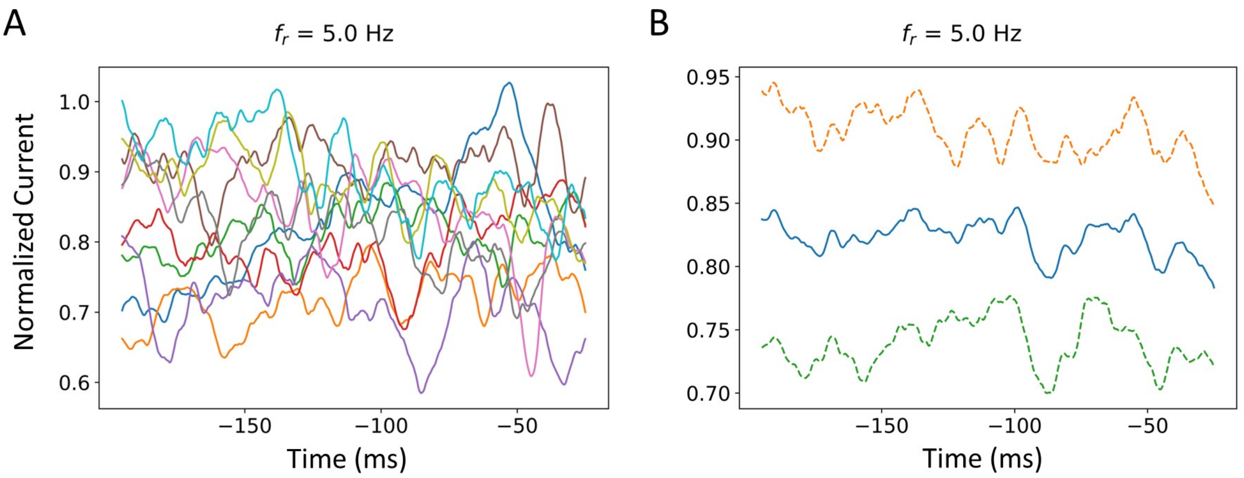


Figure S9: **Additional STA plot - IM and Ih combined.**
For $f_r$ of 5 Hz, IM and Ih are combined as described in main text Methods. **(A)** Normalized STA plots where the different colors represent different representative sets. **(B)** Blue line is the mean of normalized currents from (A). The dashed lines are $\pm$ 1 standard deviation of the normalized currents from the mean.

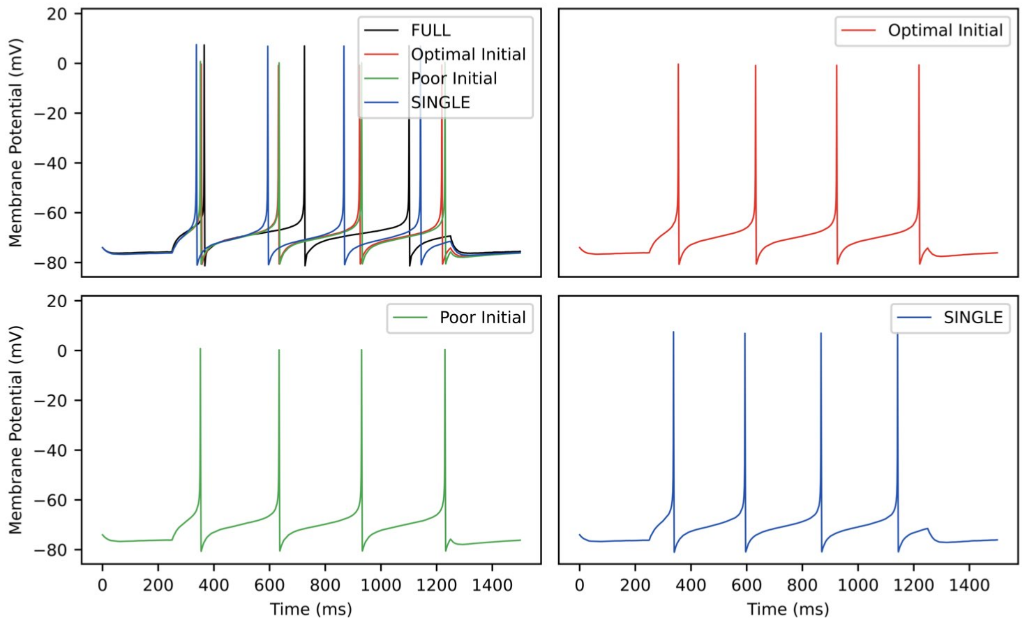


Figure S10: **RAUKF model comparisons.** Traces of FULL (black), SINGLE (blue) and RAUKF model poor and optimal estimates (green and red) to a step 60 pA step current for one second. The traces of SINGLE and RAUKF are overlayed atop FULL's in the top left. The remaining corners are the traces SINGLE and RAUKF separated for comparison.

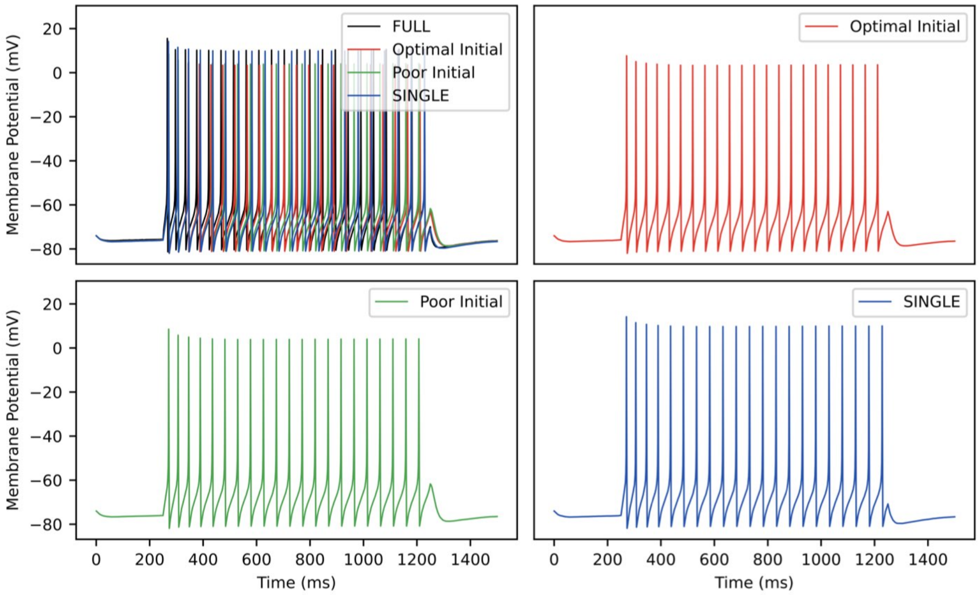


Figure S11: **RAUKF model comparisons.** Traces of FULL (black), SINGLE (blue) and RAUKF model poor and optimal estimates (green and red) to a step 60 pA step current for one second. The traces of SINGLE and RAUKF are overlayed atop FULL's in the top left. The remaining corners are the traces SINGLE and RAUKF separated for comparison.
